# Dynamic RhoA-Mediated Remodeling of Ordered Membrane Domains

**DOI:** 10.1101/2025.06.18.658998

**Authors:** Soheila Sabouri, Lucas J. Handlin, Clémence Gieré, Nicolas L.A. Dumaire, Gege Guzman, Haya Alkhateeb, Maria De Jesus Negrete, Austin D. Long, Wenwen Guo, Natalie L. Macchi, Donald W. Hilgemann, Aubin Moutal, Gucan Dai

**Author notes:** Correspondence: Gucan Dai.

## Abstract

The formation of ordered proteolipid membrane domains (OMDs) within the plasma membrane has emerged as a fundamental process that modifies membrane function, particularly in response to cell stresses that promote pathological states. Here, we identify a previously unrecognized role for the small GTPase RhoA to rapidly modulate OMDs, thereby linking cytoskeletal remodeling and membrane mechanics to OMD formation. Pharmacological and optogenetic manipulation of RhoA rapidly altered OMD dimensions in both human cell lines and dorsal root ganglion (DRG) nociceptors. The RhoA-dependent OMD expansion required actin remodeling, changes in membrane mechanical state, and protein palmitoylation. Furthermore, in a spared nerve injury (SNI) model of neuropathic pain associated with altered membrane mechanics, disrupted OMD organization, and neuronal hyperexcitability, RhoA activation promoted OMD remodeling and was accompanied by reduced HCN channel activity and action potential firing. These findings highlight alterations in plasma membrane physical properties, including changes in OMD organization, as key features of neuropathic stress. RhoA-driven OMD remodeling, mediated by diaphanous-related formin and Rho-associated kinase, may serve as a compensatory membrane adaptation that counteracts neuropathic hyperexcitability.

## Introduction

Cell membranes are laterally heterogeneous, forming dynamic lipid domains that vary in composition and size and act as platforms for localized cell signaling^1,2^. In model membranes, phase separation yields liquid-ordered (L_o_) and liquid-disordered (L_d_) phases^3,4^; In living cells, we use the term ordered membrane domains (OMDs) to describe membrane regions enriched in lipids and proteins that exhibit increased molecular order and tighter packing relative to the surrounding membrane^5–7^. While conceptually related to lipid “rafts”, OMDs are defined operationally via fluorescent and solvatochromic probes that report membrane order^5,6,8,9^, without assuming the low abundance or specific lipid composition typically implied by the raft concept. Compared with more disordered membrane regions, OMDs may differ in lipid packing, lipid-bilayer thickness, and/or protein partitioning^3,10^. In addition, OMDs undergo dynamic reorganization in dimension and clustering state^10^. Importantly, changes in membrane order and nanoscale lipid organization are implicated in many biological processes, including Alzheimer’s disease^11,12^, viral infections^13^, immune signaling^1,14^, and neuropathic pain^5,6^. Given these roles, it is essential to identify key regulators of OMD homeostasis.

RhoA, a small monomeric GTPase belonging to the Ras superfamily, has multiple well-characterized cellular functions, including regulation of actin cytoskeleton (actin stress fibers and assembly of focal adhesions), cell migration, cell shape, and endocytosis^15–18^. Guanine nucleotide exchange factors (GEFs) are responsible for the exchange of GDP for GTP and therefore activate RhoA^16,19^. Inactive, GDP-bound RhoA is predominantly cytosolic, whereas active, GTP-bound RhoA associates with the plasma membrane to engage downstream effectors. In contrast, RhoB mostly associates with endosome membranes, whereas RhoC, structurally more similar to RhoA, localizes to the plasma membrane and tends to be dysregulated during various types of cancer^20^. Importantly, RhoA activity is markedly elevated following peripheral nerve injury, which is important for nerve regeneration^21^. Cytokines, particularly those involved in neuroinflammation after nerve damage (e.g., TNF-α and IL-1β), stimulate GEFs to promote RhoA signaling in neurons^22–24^. In addition, following nerve injury, myelin-derived proteins such as myelin-associated glycoprotein (MAG) and Nogo-A, as well as signaling lipid sphingosine-1-phosphate (S1P) and lysophosphatidic acid (LPA) also activate RhoA in peripheral neurons^23,25–27^. In summary, RhoA, a membrane-associated small GTPase, is a central regulator of the cytoskeleton in sensory neurons and a mediator of the signaling cascade that regulates nerve regeneration. A less understood aspect of its function, however, is its impact on the nanoscale lipid compartmentalization and OMD organization.

Emerging evidence indicates that activation of small G proteins drives an OMD-dependent, clathrin/dynamin-independent type of endocytosis known as OMD endocytosis (OMDE) or massive endocytosis (MEND)^28–32^. Upon stimulation by various types of cell stress, the plasma membrane reorganizes to coalesce OMDs which subsequently serve as sites for initiating endocytosis^29,31,33^. Overall, OMDE is enhanced by supplementation with pyruvate, moderate concentration of palmitate/albumin complexes, and calcium transients, but is inhibited by fatty acid deprivation, treatment with 2-bromopalmitate, blockade of the mitochondrial permeability transition pore (PTP), or inhibition of palmitoyl acyltransferases (e.g., DHHC enzymes)^28–31,33^. Given the established links between OMDE, cytoskeletal remodeling, and RhoA activation^31^, we aimed to define the effects of RhoA activation on OMD properties.

Here we use fluorescence lifetime imaging microscopy (FLIM) combined with Förster resonance energy transfer (FRET) to investigate how RhoA activity influences the OMD organization. We provide evidence supporting that RhoA activation modulates OMD organization through cytoskeletal remodeling and modulation of the membrane mechanical state. Altered RhoA activity is associated with changes in HCN ion channel activity and action potential firing in sensory neurons. Furthermore, the downstream RhoA effectors diaphanous-related (mDia) formin and Rho-associated kinase (ROCK) play critical roles in regulating membrane properties. Inhibition of the HCN activity by RhoA activation and the associated dampening of membrane hyperexcitability may have evolved as an adaptation to chronic pain. These findings reveal a mechanistic link between small GTPases, the cytoskeleton, lipid domains, membrane mechanics, and excitability.

## Results

### Rho modulates OMD organization as quantified by FLIM-FRET

Cholera toxin subunit B (CTxB) conjugated FRET pairs were used as probes to determine relative changes in OMD dimension^5^. These probes are pentamers derived from *Vibrio cholerae* and have a high affinity for GM1 gangliosides in OMDs^34,35^. As previously demonstrated^5,6,36^, we used Alexa Fluor 488 (AF488) CTxB conjugates as the FRET donor and Alexa Fluor 647 (AF647) CTxB conjugates as the FRET acceptor, with a Förster critical distance (R_0_) value around 52 Å^36^. In principle, the radius of OMD determines how many CTxB probes can be accommodated within, remain in proximity to produce FRET. OMDs with a radius exceeding 10 nm have a high probability of accommodating multiple CTxB probes, whereas those with a radius smaller than 10 nm can only accommodate a single probe^5,35^. A higher FRET efficiency value is therefore indicative of a larger OMD, allowing the CTxB FRET pair to detect relative changes in OMD dimension^5,6^.

Live-cell time-resolved FLIM imaging was used to quantify FRET efficiency that reports the relative change in OMD dimension in tsA-201 cells, a HEK293T-derived cell line with an adherent morphology that facilitates membrane-localized fluorescence measurements^5,6^. The frequency-domain lifetime information of each pixel in the confocal image was transformed into a corresponding pixel on a complex plane coordinate, so-called phasor plot (Fig.1 a, b)^37^. This phasor plot displays a semicircular shape, which serves as a reference for pure single-exponential lifetimes^5,38^. Multiexponential lifetimes are found within the boundaries of this semicircle. Shorter lifetimes exhibit clockwise movements along the semicircle, indicating increased FRET^37,39–41^. With the addition of both FRET donor (CTxB-AF488) and acceptor (CTxB-AF647), there was a clockwise shift towards a shorter lifetime compared to the donor-only lifetime, indicating FRET (Fig. 1a). To calculate FRET efficiency, we used the FRET trajectory method on the phasor plot^38^. The FRET trajectory started at the zero FRET efficiency position, aligned with the donor-only cursor, and moved through the measured phasor species. As previously demonstrated and reproduced here, this CTxB-based FRET efficiency was 8.8 ± 1.1% for tsA-201 cells (Fig. 1 c, d)^5,6^.

**Fig. 1.**
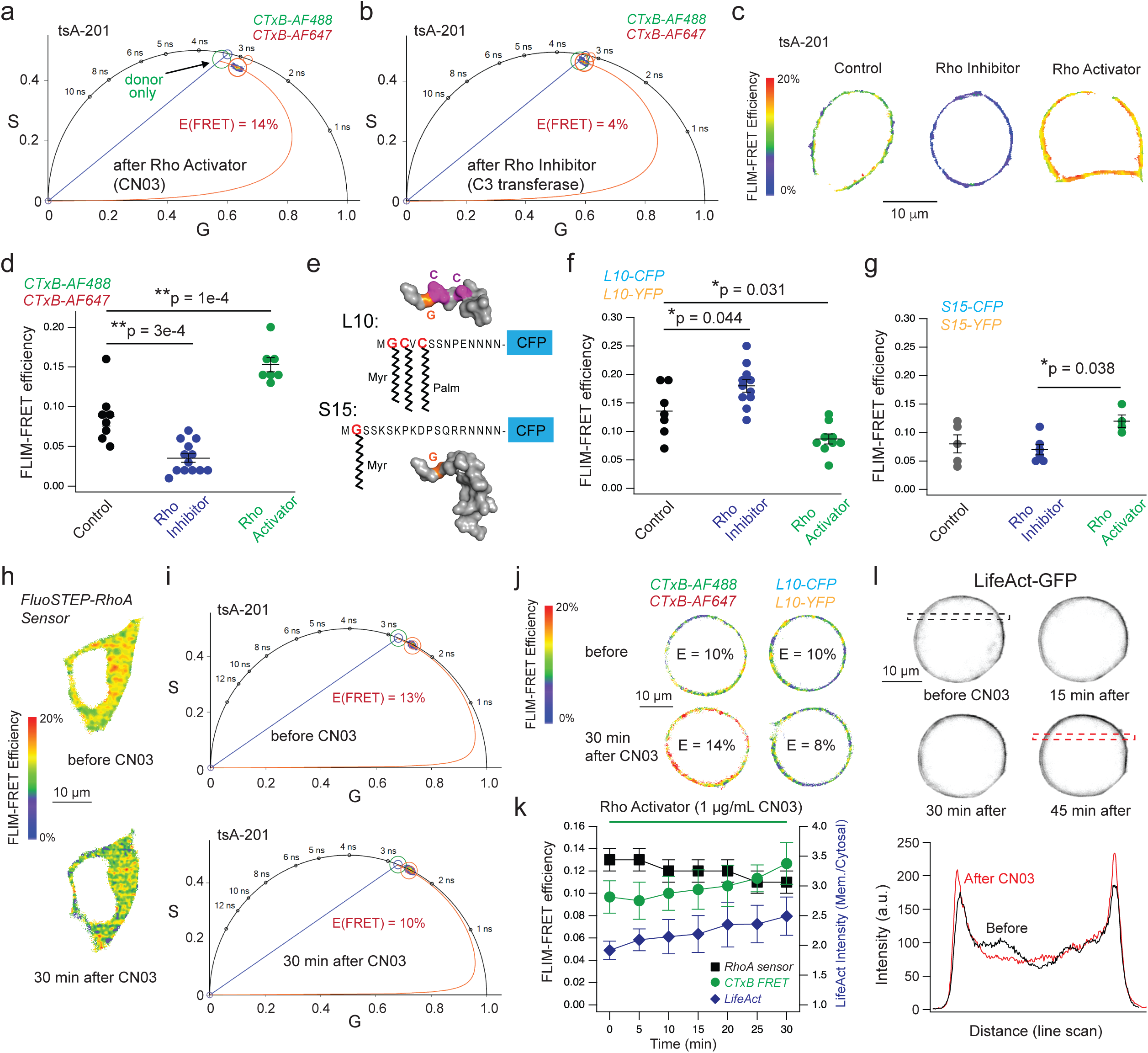
Rho activation modulates the OMD organization quantified using FLIM-FRET. **a**-**b**, Representative phasor plot of tsA-201 cells labeled by CTxB-AF488 and CTxB-AF647, without (a) and with (b) treatment of Rho inhibitor. In phasor coordinates, G and S represent the real (cosine) and imaginary (sine) components, respectively, of the normalized discrete Fourier transform of the fluorescence decay. The donor (AF-488) alone phasor is indicated as the green cursor. The trajectory curve in red begins at 0% FRET efficiency (donor only), progresses through the measured phasor, and ends at 100% efficiency, which overlaps with the background (0,0) coordinate. **c**, Representative FLIM-FRET based heatmap images of an equatorial-plane view of tsA cells, illustrating the effects of Rho inhibitor and activator on the CTxB-based FLIM-FRET efficiency. **d**, Summary of effects of Rho inhibitor and activator on the CTxB-based FLIM-FRET efficiency in tsA cells as shown in panel c, n = 8 for the untreated control, n = 13 for the Rho inhibitor, and n = 7 for the Rho activator. Data are mean ± s.e.m., **P < 0.01, one-way ANOVA. **e**, Schematic illustration of L10-and S15-CFP probes. AlphaFold 3-generated structures of L10 and S15 are also shown. **f**-**g**, Summary of analogous experiments testing the Rho inhibitor and activator using the L10-based (n = 7 – 11 cells, f) or S15-based (n = 4 – 6 cells, g) FLIM-FRET pairs in tsA cells. Data are mean ± s.e.m., *P < 0.05, one-way ANOVA. **h-i**, Representative FLIM-FRET based heatmap images (h) and phasor plots (i) of tsA cells expressing the FluoSTEP-RhoA biosensor before and after 30 min of CN03 treatment (1 µg/mL). **j**, Representative FLIM-FRET based heatmap images of the CTxB-and L10-based FRET pairs expressed in tsA cells before and after 30 min of CN03 treatment. **k**, Time course showing changes in RhoA biosensor FRET (n = 4, p = 0.016 at 30 min endpoint), CTxB-based FRET (n = 3, p = 0.035 at 30 min endpoint), and the membrane-to-cytosol fluorescence intensity ratio of LifeAct-GFP (n = 4, p = 0.046 at 30 min endpoint) following CN03 treatment, mean ± s.e.m., two-sided paired t-test. **l**, Representative LifeAct-GFP membrane fluorescence during CN03 treatment and line-scan analyses across the dashed box, highlighting the increase in membrane-associated signal.

We found that treatment with purified exoenzyme C3 transferase (1 μg/mL, > 4 hrs), a specific Rho inhibitor, significantly reduced CTxB-based FRET efficiency in tsA cells. C3 transferase, derived from *C. botulinum*, inhibits Rho GTPases through ADP-ribosylation of the effector-binding domain^42,43^. Following treatment, the FRET efficiency decreased to 3.5 ± 0.5% (Fig. 1 b-d), indicating a disruption or dispersion of OMDs. To examine the effect of Rho activation, we applied CN03 (1 μg/mL, > 4 hrs), a Rho activator derived from bacterial cytotoxic necrotizing factor (CNF) toxins that deamidates residue Q63 in the switch II region of Rho GTPases^44,45^. This modification maintains Rho in a constitutively active state^44,45^. CN03 is specific to Rho and does not modify other Ras family GTPases like Rac and Cdc42^44,45^. CN03 treatment markedly increased FRET efficiency to 15.3 ± 0.9% (Fig. 1 c, d), suggesting larger OMDs. These effects probably depended on endogenous, plasma membrane-localized RhoA (P61586; intensity: 6.43), which has significant expression in HEK293 cells, according to data from ProteomicsDB^46^. Together, these results suggest that RhoA activity modulates OMD organization in tsA cells—its activation drives domain coalescence, whereas its inhibition causes domain dispersion.

To further assess changes in OMD organization during Rho activation, we performed FLIM-FRET using the L10-CFP/L10-YFP FRET pair, which preferentially partitions into OMDs through a mechanism independent of CTxB^5,9,47^. L10 derives from the first 10 amino acids of the Lck kinase, which contains two palmitoylation sites (Fig. 1e). Unlike the CTxB-based probes, for which increased FRET reflects a greater likelihood of CTxB colocalization within OMDs, the L10-based system reports FRET through a distinct mechanism: decreased FRET reflects increased spacing between OMD-anchored L10 probes as OMDs expand^5,9^. We observed that the L10-based FRET increased following Rho inhibitor treatment and decreased with Rho activator treatment (Fig. 1f), further supporting the role of RhoA activation in OMD modulation. As an additional test, we used the FRET pair S15-CFP and S15-YFP, which localize to disordered membrane regions outside of OMDs^5,9,47^. The S15 probe contains only a single myristoylation site without palmitoylation sites (Fig. 1e). FLIM-FRET changes in the S15-based CFP/YFP pair were less pronounced following treatment with Rho activators or inhibitors, compared to the L10-based FRET changes, suggesting that Rho activity has a more indirect impact on disordered membrane regions (Fig. 1g).

### Acute RhoA activation promotes OMD and actin remodeling

To minimize cell-to-cell variability and enable real-time monitoring on a shorter timescale, we performed paired FLIM-FRET measurements in the same cell before and after CN03 treatment. We first determined whether acute CN03 exposure was sufficient to activate RhoA in tsA cells. Cells expressing a genetically encoded and FRET-based RhoA biosensor FluoSTEP-RhoA were imaged before and after CN03 application^48^. Acute treatment with CN03 (1 μg/mL, 30 min) produced a noticeable decrease in the FRET of the RhoA sensor, confirming effective activation of RhoA within 30 min under our experimental conditions (Fig. 1 h, i). Additionally, a distinct FRET-based RhoA biosensor, RhoA2G^49^, independently confirmed CN03-induced RhoA activation (Supplementary Fig. 1 a, b). We next examined whether this timescale was sufficient to alter OMD organization. Using the CTxB-based FLIM-FRET assay, we found that 30 min CN03 treatment significantly increased FRET efficiency within individual cells (Fig. 1 j, k), similar to the enhancement observed with cholesterol supplementation for this type of FRET^5,6^. Consistent with this result, L10-based FLIM-FRET decreased after CN03 treatment in the same-cell experiments, further supporting enlarged OMDs (Fig. 1 j, k and Supplementary Fig. 2d). In addition, control experiments without CN03 treatment showed minimal changes in CTxB-or L10-based FRET efficiency over the same imaging period, indicating that probe redistribution, endocytosis, or photobleaching of FRET donor at room temperature did not significantly influence the measurements (Supplementary Fig. 2 a-d). Moreover, fluorescence anisotropy measurements using a genetically encoded cholesterol sensor GRAM-W revealed no significant changes following CN03 treatment^6^, suggesting that the overall level of free cholesterol at the plasma membrane remained largely unchanged (Supplementary Fig. 2 e-f).

Because RhoA is a major regulator of actin cytoskeletal dynamics, we next investigated whether acute Rho activation was accompanied by remodeling of F-actin. Cells expressing LifeAct-GFP were imaged before and after CN03 treatment. Consistent with enhanced actin polymerization, CN03 increased LifeAct-GFP fluorescence at the cell periphery and plasma membrane, indicating an accumulation of cortical F-actin following RhoA activation (Fig. 1 k, l). The temporal correlation between RhoA activation, cortical actin enrichment, and OMD enlargement suggests that RhoA promotes membrane lipid compartmentalization through rapid reorganization of the cortical actin network.

### Optogenetic activation of RhoA rapidly promotes OMD remodeling

We used an improved Light-inducible Dimerization (iLID) system to achieve rapid and specific activation of RhoA (Fig. 2)^50–52^. This improved optogenetic system has several advantages over chemical dimerization systems and previous versions of light-induced systems, including faster onset kinetics and the ability to be turned on and off reversibly^51,53,54^. The iLID system is activated within seconds following blue light excitation and reverts to the dark state within minutes^50^, with a high recruitment efficiency and a considerable localization to the plasma membrane once activated^52,53,55^. The iLID optogenetic system involves the optical control of a RhoA-selective GEF (Fig. 2 a, b). Specifically, blue light enables the physical interaction of a protein SspB and its peptide binding target SsrA. The mCherry-tagged SspB is fused to a RhoA-selective GEF, in this case LARG^56^, and SsrA is fused to the blue light sensitive LOV2 domain (iLID-CaaX), which is targeted to the membrane via the CaaX anchor motif^50^. Upon 488 nm light excitation, the LOV2 domain, specifically the J_α_ helix, changes conformation so that SsrA becomes available for binding by SspB^57^. This allows for LARG to translocate to the membrane and activate RhoA. LARG preferentially activates RhoA over RhoB and RhoC and does not activate other Rho family proteins^58^. We confirmed blue-light induced LARG translocation through visualization of increased membrane-localized mCherry fluorescence following photoactivation (Fig. 2b). The mCherry was translocated from the cytosol to the membrane quickly, with a time constant of ∼10 seconds (Fig. 2c). RhoA activation was assessed using the localization-based biosensor mNeonGreen-3xrGBD^59^, which binds to active RhoA, as indicated by an increase in membrane-localized mNeonGreen fluorescence (Supplementary Fig. 1 c, d).

**Fig. 2.**
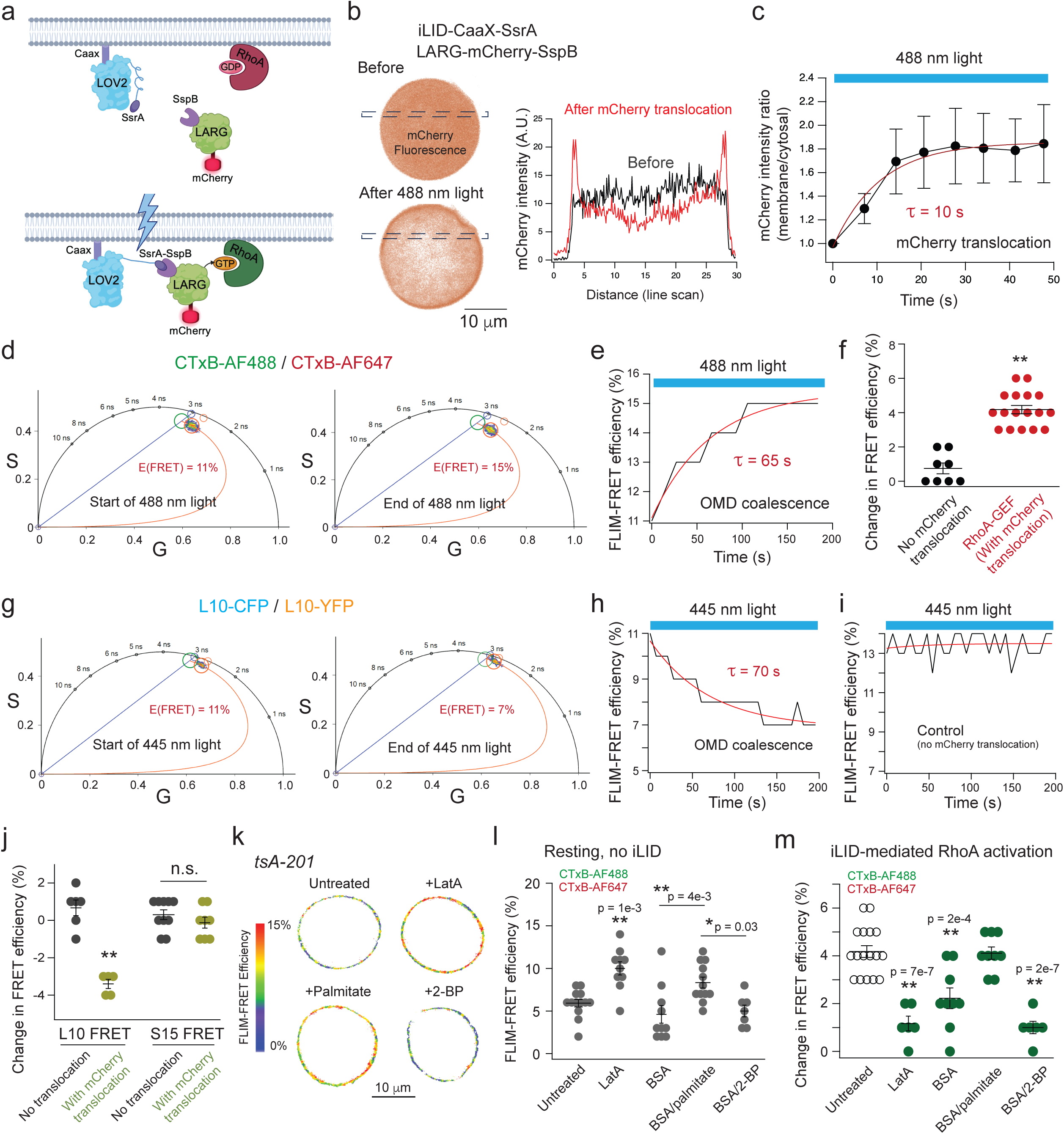
Activation of RhoA using a light-inducible dimerization (iLID) system. **a**, Schematic of the iLID module before and after 488-nm illumination. Photoexcitation triggers a conformational change in LOV2–CaaX–SsrA that exposes the SsrA motif and enables SspB binding. **b**, Intensity-based heatmaps of tsA-201 cells before and after illumination show that activation of the iLID system drives rapid redistribution of LARG–mCherry from the cytosol to the plasma membrane. Line-scan profiles confirm increased membrane-associated fluorescence concurrent with reduced cytosolic signal. **c**, Time course of LARG–mCherry recruitment quantified as the ratio of membrane-localized to cytosolic fluorescence (n = 4 cells). **d**, Representative phasor plots from cells labeled with CTxB-AF488 and CTxB-AF647 and expressing the iLID module. Comparison of the start and the end of 488-nm excitation reveals an increase in FRET efficiency. **e**, Corresponding time course of FLIM-FRET efficiency between CTxB-AF488 and CTxB-AF647. **f**, Summary of CTxB-based FRET changes in cells exhibiting clear LARG–mCherry translocation (n = 17) versus those lacking detectable translocation (n = 8), mean ± s.e.m., **P = 8e-7, two-sided t-test. **g**, Representative phasor plots from cells expressing L10-CFP/YFP and the iLID module. **h**-**i**, Time course of L10-based FRET during 445-nm illumination in cells with (h) or without (i) LARG–mCherry membrane recruitment, demonstrating that iLID-evoked changes in FRET are coupled to RhoA activation. **j**, Summary of L10-and S15-based FRET responses in cells with or without LARG–mCherry translocation (L10: n = 6 without, n = 5 with; S15: n = 10 without, n = 8 with; **P = 1e-7, one-way ANOVA). **k**, Representative FLIM–FRET heatmaps of CTxB-labeled tsA cells under control conditions or following treatment with Lat-A, palmitate, or 2-BP, illustrating how cytoskeletal disruption or altered palmitoylation modulates baseline FRET efficiency. **l**, Quantification of baseline CTxB-based FRET efficiencies for control (n = 13), Lat-A (n = 11), BSA (n = 10), palmitate (n = 12), and 2-BP (n = 7) conditions (two-sided t-test). **m**, Summary of iLID-evoked changes in CTxB-based FRET across the same pharmacological conditions (control: n = 17, Lat-A: n = 6, BSA: n = 9, palmitate: n = 9, 2-BP: n = 6; **P < 0.01, one-way ANOVA).

FLIM-FRET between CTxB-AF488 and CTxB-AF647 was measured during global photoactivation of the cell. In this protocol, the 488 nm laser served a dual purpose: activating the optogenetic system and exciting the FRET donor CTxB-AF488. We found the FRET efficiency significantly increased following endogenous RhoA activation, on average around 4% (Fig. 2 d-f). There was an exponential increase that had a time constant of 88 ± 6 seconds (n = 14, Fig. 2e). Control experiments performed in the absence of the FRET acceptor, but in the presence of the optogenetic system, revealed no change in the FRET donor (CTxB-AF488) lifetime upon light activation (Supplementary Fig. 3). In addition, control cells lacking LARG-mCherry-SspB (untransfected) or iLID-CaaX (undetectable mCherry translocation upon photoactivation) exhibited only an average increase of 1% in FRET efficiency (Fig. 2f and Supplementary Fig. 4 a-d, 4a). These results indicate that activation of endogenous RhoA in tsA201 cells promotes the expansion of OMDs. Throughout these measurements, cell morphology was monitored, and no appreciable changes in cell shape were observed despite the alterations in FLIM-FRET signals (Supplementary Fig. 5a).

A key advantage of the iLID optogenetic system is its ability to rapidly and reversibly manipulate RhoA activity. To determine whether OMD remodeling is likewise reversible, we implemented an activation–deactivation paradigm. Illumination with 488 nm light rapidly activates RhoA, increasing CTxB-based FRET, consistent with OMD enlargement (Supplementary Fig. 4e). After illumination ceased, LARG-mCherry returned to its cytosolic localization within 5 min, whereas the CTxB-based FRET signal recovered more gradually, reaching near-baseline levels in approximately 20 min. A second round of illumination elicited a comparable increase in CTxB-based FRET (Supplementary Fig. 4 e-g), demonstrating that RhoA-induced OMD remodeling is reversible as RhoA activity declines. Together, these findings provide evidence that OMD remodeling is reversibly and bidirectionally regulated by RhoA activity.

We also tested the iLID system and its effects on OMDs using the L10-based FRET pairs in tsA cells. Like the CTxB experiments, photoactivation was applied using 445 nm light, which simultaneously activated the optogenetic system and excited the FRET donor L10-CFP. Upon RhoA activation, we observed a considerable 3.4 ± 0.2% decrease (n = 5) in FRET efficiency between coexpressed L10-CFP and L10-YFP, with a time constant comparable to the kinetics of the FRET increase observed for CTxB-based FRET pairs (Fig. 2 g-j). Control cells lacking LARG-mCherry-SspB translocation showed little changes in FRET (Fig. 2i). In tsA cells expressing the optogenetic system and S15-CFP and S15-YFP FRET pairs, photoactivation induced no measurable change in S15-based FRET efficiency (change in FLIM-FRET efficiency = 0 ± 0.3%), indicating that the effects observed with the L10 pair are specific to changes in lipid-ordered domains (Fig. 2j). In summary, these complementary results support the conclusion that RhoA activation drives the rapid coalescence of OMDs.

### Dependence of RhoA-driven OMD remodeling on cytoskeleton and protein palmitoylation

Considering the prominent role of RhoA in cytoskeletal remodeling, we tested whether disrupting the actin cytoskeleton would affect OMDs and the action of RhoA. Treatment with latrunculin A (Lat-A, 5 μM, 10 min), a potent actin polymerization inhibitor^60^, increased the baseline CTxB-based FRET efficiency in tsA cells but nearly abolished the RhoA-induced increase in FRET following optogenetic activation (Fig. 2 k-m). These results suggest that an intact actin cytoskeletal network is required to maintain OMD organization. Disruption of cytoskeletal support results in larger OMDs, thereby impairing the ability of RhoA to further promote OMD expansion (Fig. 2l).

Given the critical role of palmitoylation in OMD formation and related endocytosis functions^61–63^, we tested whether increasing cellular palmitate levels and altering protein palmitoylation modulate RhoA’s effects on OMDs. To achieve this, tsA cells were treated with palmitate chelated to bovine serum albumin (palmitate/BSA 6:1; 50 μM, 20 min), and control cells were treated with BSA alone under identical conditions. Palmitate/BSA treatment increased the baseline CTxB-based FRET efficiency compared to BSA controls, indicating enlargement of OMDs (Fig. 2 k-l). In contrast, BSA alone slightly reduced baseline FRET efficiency, likely due to its scavenging effect on free fatty acids present in the culture medium (Fig. 2 k-l). These results suggest that incorporation of saturated fatty acids into the plasma membrane promotes the formation or stabilization of larger OMDs. In the optogenetic RhoA activation experiment, BSA treatment attenuated the RhoA-induced FRET increase, whereas palmitate/BSA treatment preserved the FRET response (Fig. 2m). Furthermore, pre-treatment with 2-bromopalmitate (2-BP; 100 μM, 1 h), a broad-spectrum inhibitor of protein palmitoylation, did not alter baseline FRET efficiency but abolished the RhoA-induced FRET increase (Fig. 2 k-m). These results indicate that protein palmitoylation is required for RhoA-dependent OMD expansion.

### RhoA activation modulates OMDs and is accompanied by a reduction in membrane excitability in nociceptor DRG neurons

We extended these findings to rat primary small-diameter dorsal root ganglion (DRG) neurons, most of which are nociceptors responsible for detecting noxious stimuli and are sensitive to changes in membrane composition^64,65^. These small DRG neurons are important for processing pain signals and possess larger OMDs than large-diameter DRG neurons and tsA cells^5^. The FRET efficiency in naïve small DRG neurons (control) was measured at 19.5 ± 2.0% (Fig. 3 a, b). For these neurons, prolonged exposure to C3 transferase (1 μg/mL, >4 hrs) led to a significant decrease in CTxB-based FRET efficiency to 10.7 ± 0.6%, consistent with shrunken OMDs (Fig. 3 a, b). Conversely, activating Rho signaling using CN03 failed to elevate FRET efficiency significantly (measured as 20.6 ± 1.6%). This suggests that OMDs in naïve nociceptors are already large, rendering further Rho-induced expansion undetectable by the CTxB-based FRET method. Then we measured the spontaneous action potential firing of naïve nociceptor DRG neurons (Fig. 3c). In control neurons, 40% exhibited spontaneous firing (n = 15), consistent with our previous report^5^. Spontaneous firing increased to 93% in Rho inhibitor-treated neurons (n = 15), whereas 45% of Rho activator-treated neurons exhibited firing (n = 11). We also tested the influence of Rho inhibition and activation on the input resistance of naïve neurons and found no significant change (Supplementary Fig. 6). These results suggest that, as in tsA cells, decreased RhoA activity reduces OMDs in naïve sensory neurons, likely modulating their excitability and signaling properties.

**Fig. 3.**
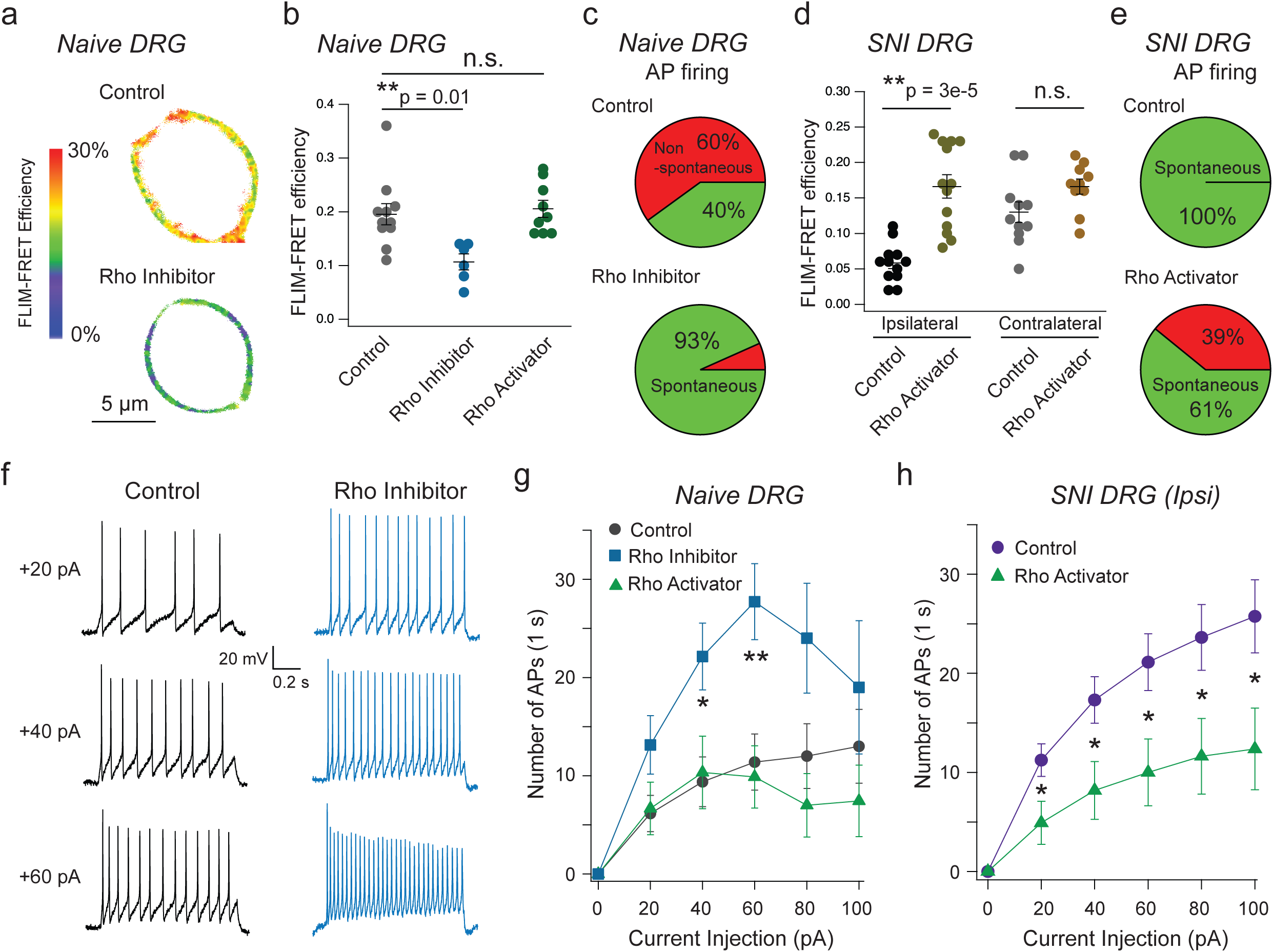
Rho-dependent modulation of OMD and associated changes in excitability in small DRG neurons. **a**, Representative FLIM–FRET heatmaps of naïve small-diameter DRG neurons labeled with CTxB illustrate the reduction in FRET efficiency reporting the change in OMD dimension following acute Rho inhibition. **b**, Quantification of CTxB-based FLIM–FRET efficiencies in naïve neurons under control conditions (n = 11 cells), after Rho inhibition (n = 6), or after Rho activation (n = 9) (mean ± s.e.m.; one-way ANOVA). **c**, Pie charts showing the proportion of naïve DRG neurons exhibiting spontaneous action potential (AP) firing in control versus Rho inhibitor–treated groups (green, spontaneous APs; red, silent). **d**, Summary of CTxB-based FLIM–FRET efficiencies in small DRG neurons from spared nerve injury (SNI) rats. Ipsilateral neurons showed increased FRET after Rho activation (control: n = 9 cells; Rho activator: n = 13). Corresponding data for contralateral neurons are shown (control: n = 11 cells; Rho activator: n = 10) (one-way ANOVA, p = 2e-6 for the ipsilateral neurons and p = 0.27 for contralateral neurons). **e**, Pie charts illustrating the fraction of ipsilateral SNI neurons that fired spontaneous APs under control conditions versus after Rho activation. **f**, Representative traces of current injection–evoked AP firing in naïve nociceptors under control conditions and after Rho inhibition, highlighting enhanced excitability with suppressed RhoA activity. **g**, Summary of current injection-elicited AP firing in naïve DRG neurons under control conditions (n = 13 patches), with Rho inhibition (n = 7), or with Rho activation (n = 9). AP counts are plotted against injected current steps (mean ± s.e.m., p = 0.02 at 40 pA and p = 0.005 at 60 pA; one-way ANOVA). **h**, Summary of current injection-evoked firing from ipsilateral SNI nociceptors under control conditions (n = 16 patches) and after overnight CN03 treatment (n = 11), two-sided t-test, p = 0.03, 0.02, 0.02, 0.03, 0.02 at 20, 40, 60, 80, 100 pA current injections; *p < 0.05, **p < 0.01.

We next tested whether activating RhoA could restore the OMD dimension in nociceptor DRG neurons isolated from rats subjected to spared nerve injury (SNI), a well-established model of neuropathic pain^66^. Small-diameter neurons from SNI animals exhibited significantly reduced CTxB-based FRET efficiency (5.9 ± 1.1%) (Fig. 3d), reflecting a disruption of OMDs typically associated with chronic injury and membrane hyperexcitability^5^. Overnight treatment with Rho activator CN03 reversed this reduction, restoring FRET efficiency (16.6 ± 1.6%) toward levels observed in naïve and contralateral SNI neuron controls (Fig. 3d). Consistently, acute treatment with CN03 significantly increased the CTxB-based FRET efficiency in the same-neuron experiment (Supplementary Fig. 5 b, c). These findings suggest that Rho signaling plays an essential role in preserving OMD architecture under pathological conditions and may offer a mechanism to stabilize membrane nanodomain organization in sensory neurons affected by nerve injury. Notably, this restoration of OMD coincided with reduced spontaneous firing: 61% of RhoA-activated SNI neurons exhibited spontaneous activity (n = 23), compared to 100% in hyperexcitable SNI controls (n = 18), consistent with our previous findings^5^ (Fig. 3e). Additionally, current injection–evoked action potential firing was potentiated by Rho inhibition but showed minimal change upon Rho activation in naïve neurons (Fig. 3f). In contrast, treatment with the Rho activator markedly reduced current injection–evoked action potential firing in ipsilateral SNI neurons (Fig. 3h). These distinct effects are consistent with smaller OMDs in hyperexcitable pain states and with RhoA activation dampening neuronal hyperexcitability in our in vitro experiments using primary cultured neurons.

To assess how the actin cytoskeleton and protein palmitoylation regulate OMD organization and neuronal excitability in naïve and SNI nociceptors, we performed CTxB-based FRET measurements analogous to those used in tsA cells (Supplementary Fig. 7 a, b). Disruption of the F-actin network with Lat-A rescued shrunken OMDs observed in SNI neurons, while having minimal effect on naïve neurons. Manipulation of palmitoylation, either by palmitate supplementation or inhibition with 2-BP, altered the FRET, consistent with the idea that protein palmitoylation is required to maintain OMD integrity. Functionally, 2-BP treatment decreased the FRET efficiency in naïve DRG neurons and was sufficient to induce neuronal hyperexcitability (Supplementary Fig. 7 a, c). In contrast, 2-BP had little additional effect on SNI neurons, which already display both hyperexcitability and shrunken OMDs (Supplementary Fig. 7 b, d). In ipsilateral SNI neurons, Rho activation failed to reduce elicited action potential firing in the presence of a subsequent Lat-A treatment, indicating that RhoA’s effect depends on an intact cytoskeletal network (Supplementary Fig. 7d). Together, these results indicate that disruption of palmitoylation-dependent OMD organization is sufficient to induce hyperexcitability in naïve neurons, whereas remodeling of the actin cytoskeleton likely mediates the OMD restoration in SNI neurons.

### RhoA regulates OMDs through cytoskeleton-dependent modulation of membrane mechanics

To examine how RhoA-induced changes of OMDs are dependent on mechanical properties of the plasma membrane, we used the fluorescent probe Flipper-TR, which reports lipid packing and lateral membrane tension through changes in its fluorescence lifetime^67–69^ (Fig. 4a). Because actin cytoskeletal dynamics are closely coupled to membrane mechanics, this approach allows us to assess how RhoA-mediated cytoskeletal remodeling influences this property of the membrane. Changes in membrane packing modulate the conformation of the dithienothiophene moieties of Flipper-TR, altering intramolecular rotation and thereby affecting fluorescence lifetime^67,68^. Thus, phasor FLIM measurements of Flipper-TR provide a nanoscale readout of changes in the local membrane mechanical environment (Fig. 4b). As a proof-of-principle experiment, we acutely applied extracellular solutions (< 3 min) with varying osmolarity to tsA cells labeled with Flipper-TR. The phase lifetime (τϕ) of Flipper increased from 4.5 ± 0.1 ns in hypertonic solution (493 mOsm) to 4.8 ± 0.05 ns in isotonic solution (326 mOsm), and to 5.2 ± 0.1 ns in hypotonic solution (154 mOsm) (Fig. 4c). Similar osmolarity-dependent changes in Flipper-TR lifetime were observed in naïve DRG neurons (Fig. 4d), consistent with the probe’s sensitivity to changes in membrane mechanics.

**Fig. 4.**
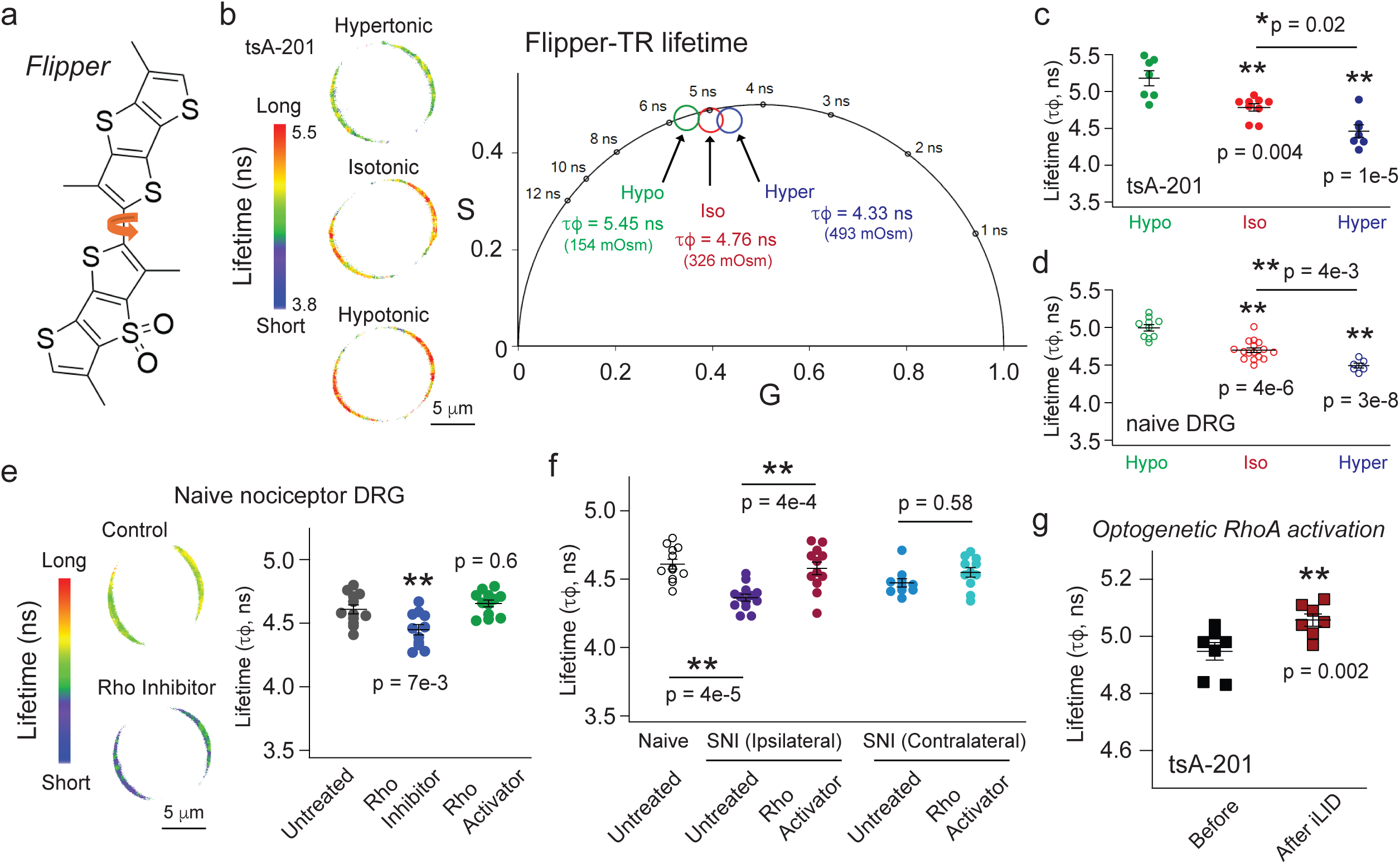
RhoA-dependent modulation of membrane mechanics measured by the sensor Flipper-TR. **a**, Schematic of Flipper-TR, a mechanosensitive probe whose fluorescence lifetime is sensitive to the local membrane mechanical environment, including lipid packing and membrane tension. Changes in membrane mechanical state alter the conformation of the dithienothiophene moieties, affecting intramolecular rotation and thereby modulating fluorescence lifetime. Flipper-TR has an additional head group (not shown) for plasma membrane localization. **b**, Representative Flipper-TR lifetime-based heatmap images of tsA cells exposed to extracellular solutions of varying osmolarity, showing graded increases from hypertonic to isotonic and hypotonic conditions, as well as the lifetime shift shown in the phasor plot. **c**-**d**, Summary of osmolarity-dependent lifetime changes in tsA cells (c) and naïve DRG neurons (d). n = 7-9 cells for tsA and n = 6-16 for naïve DRG, mean ± s.e.m., one-way ANOVA. **e**, Pharmacological manipulation of RhoA activity in naïve DRG neurons revealed that Rho inhibition lowered Flipper-TR lifetime, whereas Rho activation produced minimal effects, n = 12 for the control and Rho inhibitor; n = 13 for Rho activator, mean ± s.e.m., one-way ANOVA. **f**, Summary data of SNI DRG neurons that exhibited reduced baseline Flipper lifetime. RhoA activation substantially increased lifetime, restoring values toward those of contralateral neurons, n = 13 for the control (ipsilateral), n = 12 for Rho activator (ipsilateral); n = 10 for the control (contralateral), n = 11 for Rho activator (contralateral), mean ± s.e.m., one-way ANOVA. **g**, Optogenetic activation of RhoA increased Flipper-TR lifetime during 488-nm illumination, n = 7, paired t-test. *p < 0.05, **p < 0.01.

Using FLIM imaging of Flipper-TR, we monitored changes in membrane mechanics during pharmacological and optogenetic activation of RhoA to test whether potential changes in lipid packing, lateral tension, and membrane mechanics accompany RhoA-induced reorganization of OMDs (Fig. 4 and Supplementary Table 1). We found that, in naïve DRG neurons, treatment with a Rho inhibitor reduced the fluorescence lifetime of Flipper-TR, whereas treatment with a Rho activator produced little effect, suggesting that the membrane mechanical environment and OMD organization were already elevated under basal conditions (Fig. 4e). Importantly, ipsilateral SNI DRG neurons exhibited a shorter baseline Flipper lifetime than either naïve or contralateral SNI neurons (Fig. 4f), indicating an altered membrane mechanical state. Treatment with a RhoA activator increased Flipper lifetime in ipsilateral neurons to levels comparable to those observed in naïve and contralateral SNI neurons (Fig. 4f). Furthermore, optogenetic activation of RhoA in tsA-201 cells substantially increased Flipper lifetime (Fig. 4g), supporting a role for RhoA in regulating the local membrane mechanical environment.

Consistent with the changes observed in OMD organization, both the actin cytoskeleton and protein palmitoylation influenced the membrane mechanical environment reported by Flipper-TR (Supplementary Fig. 8a). In naïve DRG neurons, 2-BP reduced Flipper lifetime, whereas disruption of the actin cytoskeleton with Lat-A had little effect. In contrast, in ipsilateral nociceptors from SNI rats, Lat-A increased Flipper lifetime, whereas 2-BP failed to further reduce it, consistent with differences in the baseline membrane mechanical state between naïve and SNI neurons. One possible explanation is that the cortical actin network normally maintains membrane reservoirs that buffer mechanical tension; thus, Lat-A-induced actin depolymerization reduces membrane reserve capacity and increases effective membrane tension^70^. Furthermore, the increase in Flipper lifetime induced by RhoA activation was abolished by Lat-A-mediated actin disruption, indicating that an intact actin network is required for RhoA-dependent regulation of membrane mechanics (Supplementary Fig. 8a). In contrast, 2-BP did not prevent the RhoA activator-induced increase in Flipper lifetime in ipsilateral SNI neurons, suggesting that palmitoylation acts upstream of RhoA signaling rather than directly mediating this response.

Moreover, we assessed the effects of the chemotherapeutic agent paclitaxel, which has previously been shown to disrupt OMD organization^5^. Because paclitaxel stabilizes microtubules, we hypothesized that its effects on membrane mechanics would differ from those of Lat-A-mediated actin disruption. Overnight treatment with 1 μM paclitaxel reduced Flipper lifetime in naïve DRG neurons, indicating an altered membrane lipid environment. In contrast, paclitaxel had little effect in ipsilateral SNI neurons, where the Flipper lifetime was already reduced (Supplementary Fig. 8b). Together, these findings suggest that actin and microtubule perturbations exert distinct effects on membrane nanodomain organization and likely on tension-related mechanics. Finally, cholesterol depletion with methyl-β-cyclodextrin (MβCD) reduced Flipper lifetime in naïve DRG neurons (Supplementary Fig. 8c), consistent with a role for cholesterol in maintaining membrane order and mechanical properties.

### Diaphanous-related formin (DRF) and ROCK contribute to the modulation of OMDs

Diaphanous-related (mDia) formin and Rho-associated coiled-coil-containing protein kinase (ROCK) are major downstream effectors of RhoA that regulates actomyosin contractility, linear actin organization, and membrane mechanical properties^71,72^ (Fig. 5a). We used the iLID optogenetic system to acutely activate RhoA and found that activation of mDia with IMM01 (10 μM) increased basal CTxB-based FRET before activating iLID but attenuated the subsequent iLID-mediated FRET increase (Fig. 5 b, c). Inhibition of mDia and ROCK with a formin inhibitor SMIFH2 (10 μM) and Y-27632 (20 μM) also reduced the iLID-mediated FRET increase, respectively (Fig. 5c). To further test whether ROCK contributes to RhoA-dependent OMD expansion, we inhibited ROCK using Y-27632 after activating RhoA with CN03 treatment. In these paired same-cell measurements, Y-27632 increased the L10-based FLIM-FRET and reduced the CTxB-based FRET efficiency in tsA-201 cells, consistent with inhibition of ROCK downstream of CN03-induced RhoA activation (Supplementary Fig. 9 a, b).

**Fig. 5.**
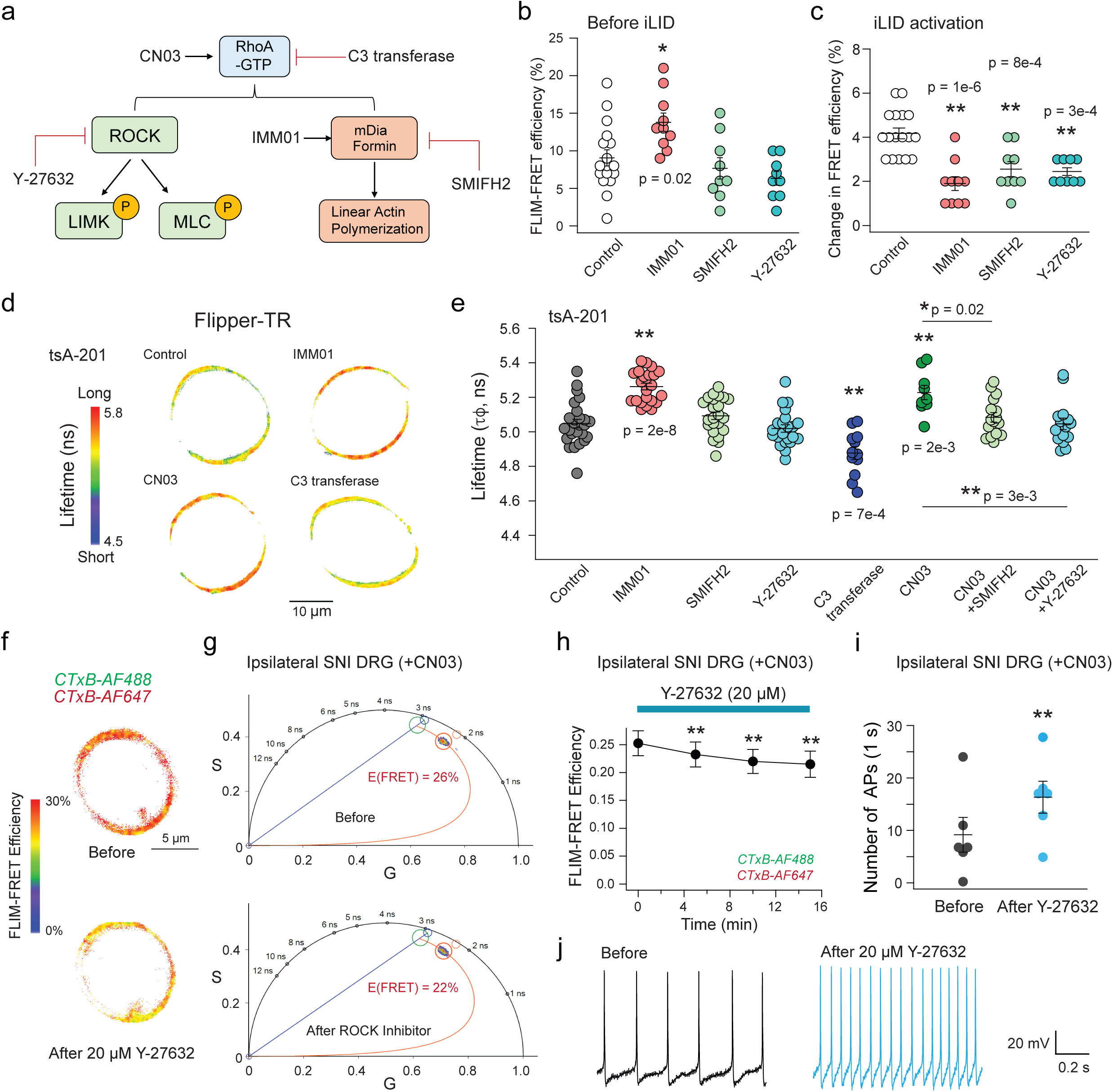
Both mDia and ROCK contribute to Rho-mediated effects. **a**, Schematic showing the downstream effectors of RhoA, including ROCK and mDia formins, and the pharmacological activators and inhibitors used in this study. **b**-**c**, Summary of the effects of IMM01, SMIFH2 and Y-27632 on CTxB-based FRET efficiency under resting conditions (start, b) or following light activation of the iLID system (c). Data are mean ± s.e.m.; one-way ANOVA; n = 17 cells for control, n = 10 for IMM01, and n = 9 for SMIFH2 and Y-27632. **d**, Representative Flipper-TR lifetime-based heatmap images of tsA cells under control conditions or following treatment with IMM01, SMIFH2, Y-27632, C3 transferase or CN03. **e**, Summary of Flipper-TR fluorescence lifetimes under the indicated conditions. Data are mean ± s.e.m.; one-way ANOVA; n = 26 cells for control, n = 23 for IMM01, n = 23 for SMIFH2, n = 22 for Y-27632, n = 12 for C3 transferase, n = 9 for CN03, n = 16 for CN03 + SMIFH2, and n = 15 for CN03 + Y-27632. **f**-**g**, Representative FLIM-FRET heatmap images (f) and corresponding phasor plots (g) showing the effect of acute Y-27632 treatment (20 μM) on CTxB-based FRET in ipsilateral SNI nociceptor DRG neurons following overnight treatment with CN03 (1 μg/mL). **h**, Summary time course of the experiment shown in f and g following acute Y-27632 treatment. Data are mean ± s.e.m.; n = 4; p = 0.001, 0.0006 and < 0.0001 at 5, 10 and 15 min, respectively. **i**, Summary of spontaneous action potential firing in ipsilateral SNI DRG neurons before and after acute Y-27632 treatment. n = 6; paired two-tailed t-test; p = 0.007. **j**, Representative recordings of spontaneous action potential firing from ipsilateral SNI DRG neurons before and after acute Y-27632 treatment following overnight incubation with CN03 (1 μg/mL).

Aligned with these findings, IMM01 increased the Flipper lifetime, whereas neither SMIFH2 nor Y-27632 significantly reduced the Flipper lifetime under basal conditions (Fig. 5 d, e). This may reflect the relatively low level of endogenous mDia or ROCK activation in tsA cells, such that inhibition of its downstream effectors has limited effects under basal conditions. However, following overnight treatment with CN03 to activate RhoA and elevate the Flipper lifetime, subsequent inhibition of either mDia or ROCK significantly reduced the Flipper lifetime (Fig. 5 d, e). Together, these results suggest that both mDia-and ROCK-dependent pathways contribute to RhoA-mediated regulation of OMD organization and membrane lipid packing in tsA-201 cells. Given that mDia promotes F-actin polymerization, these findings suggest that the mDia-actin pathway plays a major role in RhoA-mediated regulation of membrane mechanics.

We next examined whether ROCK and mDia contribute to membrane properties of DRG neurons. Again, to minimize cell-to-cell variability, we performed acute same-cell measurements before and after the application of Y-27632. Pharmacological inhibition of ROCK using Y-27632 (20 μM) reduced CTxB-based FRET in naïve nociceptor DRG neurons (Supplementary Fig. 9 c, d). In addition, same-cell patch-clamp recordings enabled direct comparison of spontaneous neuronal firing before and after treatment, revealing that spontaneous activity increased in naïve neurons after ∼5 min of Y-27632 treatment (Supplementary Fig. 9 e, f). These results indicate that ROCK activity is required for maintaining normal OMDs and membrane excitability for naïve nociceptor neurons. We next tested whether ROCK regulates the RhoA-dependent expansion of OMDs in SNI DRG neurons. Similar same-cell FLIM-FRET experiments in ipsilateral DRG neurons showed that Y-27632 decreased CTxB-based FRET within 10-15 min (Fig. 5 f-h), suggesting that ROCK activity facilitates enlarged OMDs in neuropathic neurons after the overnight CN03 treatment. However, Y-27632 only partially reversed the CN03-induced changes, suggesting that other downstream effectors, such as mDia, also contribute to RhoA-dependent OMD expansion. Furthermore, we examined the effect of ROCK inhibition on neuronal firing. In same-cell experiments using ipsilateral SNI neurons, acute application of Y-27632 significantly increased spontaneous action potential firing (Fig. 5 i, j). These findings indicate ROCK as a downstream effector of RhoA-mediated OMD expansion and suggest that ROCK-dependent membrane organization contributes to the suppression of spontaneous activity in neuropathic nociceptors. Furthermore, we found mDia inhibitor SMIFH2 slightly decreased the CTxB-based FRET whereas mDia activator IMM01 increased the FRET in ipsilateral DRG neurons (Supplementary Fig. 10 a, b). Consistently, application of formin inhibitor SMIFH2 (10 μM) increased the action potential firing in naïve DRG neurons (Supplementary Fig. 10 c, d). Collectively, similar to HEK cells, both ROCK and mDia may contribute to the RhoA-mediated OMD organization in small nociceptor DRG neurons.

### The effect of hypotonicity-induced swelling on OMD organization

To test whether acute changes in membrane mechanics are sufficient to promote OMD reorganization, we treated tsA cells and DRG neurons with acute hypotonic stress to induce cell swelling and membrane stretching. For the same-cell experiments, cells initially maintained in isotonic solution were exposed to a 1:1 mixture of isotonic (326 mOsm) and hypotonic (154 mOsm) solutions, resulting in a final osmolarity of 240 mOsm. This treatment increased CTxB-based FRET by ∼2% within 4 min (Fig. 6 a, b). The increase in CTxB-FRET is consistent with increased co-occupancy of CTxB molecules within OMDs, as expected if elevated membrane stretching/tension promotes OMD coalescence^73^. In agreement with enlarged OMDs, L10-CFP/L10-YFP FRET efficiency decreased significantly under hypotonic conditions, as assessed using same-cell measurements before and after treatment (Fig. 6 c). In separate-cell experiments, we applied a stronger hypotonic treatment (154 mOsm for ∼3 min), which similarly decreased the L10-based FRET (Fig. 6d) and increased the CTxB-based FRET efficiency (Fig. 6e). This effect persisted in the presence of overnight treatment of RhoA inhibitor C3 transferase (Fig. 6f), suggesting that acute hypotonicity-induced OMD remodeling does not require RhoA activation.

**Fig. 6.**
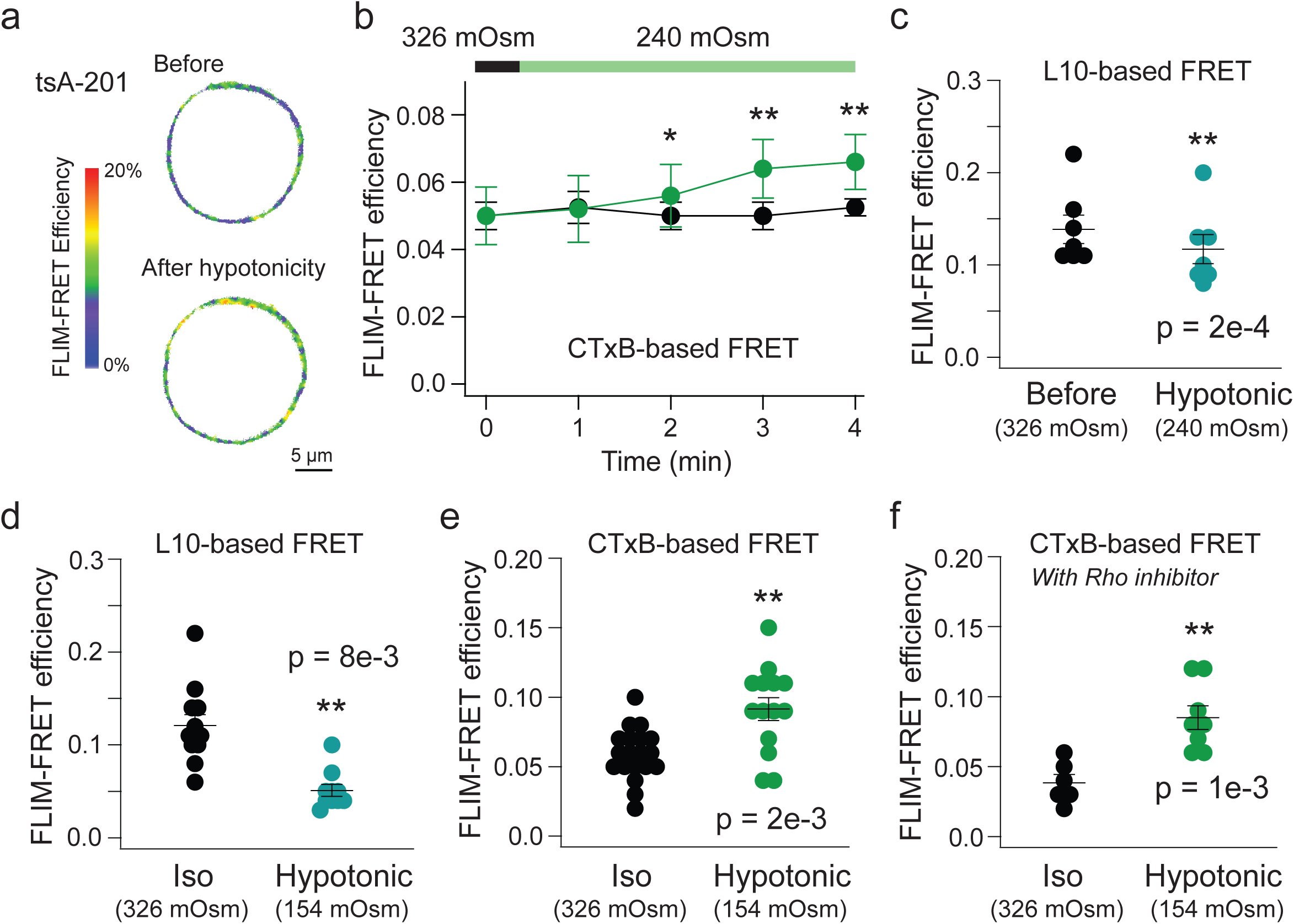
Effect of hypotonicity on FRET probes reporting OMD organization. **a**, Representative FLIM-FRET based heatmap images of the CTxB-based FRET pairs expressed in tsA cells during the hypotonic treatment (240 mOsm), with the initial 326 mOsm isotonic condition shown before treatment. **b**, Summary time course of the CTxB-based FRET efficiency when changing the bath solution from 326 mOsm to 240 mOsm; the untreated control showed no change in FRET. Data are mean ± s.e.m.; n = 6 with the treatment and n = 4 for untreated controls, p = 0.1, 0.04, 0.002, 6e-4 at 1, 2, 3, 4 min time points, respectively, paired two-tailed t-test. **c**, A similar same-cell experiment applying 240 mOsm hypotonic swelling as in panel b, but using the L10-based FRET. Data are mean ± s.e.m.; n = 7, paired two-sided t-test. **d**, Acute (∼3 min) hypotonic treatment at 154 mOsm decreased the L10-based FRET. Mean ± s.e.m., n = 12 cells for the control, n = 10 cells for hypotonic condition, two-sided student’s t-test. **e**-**f**, Acute (∼3 min) hypotonic treatment at 154 mOsm increased the FLIM-FRET efficiency of tsA-201 assessed using the CTxB-based FRET pair CTxB-AF488 and CTxB-AF647, without (e) or with (f) overnight treatment with Rho inhibitor (C3 transferase). Mean ± s.e.m., n = 19 cells for the control, n = 14 cells for hypotonic condition in panel e and n = 6 cells for the control, n = 8 cells for hypotonic condition in panel f, *p < 0.05, **p < 0.01 using two-sided student’s t-test.

In naïve DRG nociceptors, which exhibit relatively high baseline FRET, the acute hypotonic treatment increased FRET efficiency only following Rho inhibition (Supplementary Fig. 11a). In SNI DRG neurons, hypotonic treatment increased FRET in ipsilateral neurons but produced no significant further increase in contralateral neurons, which exhibited relatively high baseline FRET (Supplementary Fig. 11b). Together, these findings suggest that the mechanical state of the membrane contributes to the RhoA-dependent OMD remodeling.

To examine how cortical F-actin responds to distinct perturbations of membrane mechanics, we imaged LifeAct-GFP in tsA cells following acute hypotonic treatment or latrunculin A (Lat-A) application (Supplementary Fig. 12). Hypotonic swelling rapidly reduced membrane-associated LifeAct-GFP fluorescence, consistent with remodeling of the cortical F-actin network, whereas Lat-A reduced peripheral LifeAct-GFP signal as expected from actin depolymerization. Although these perturbations alter the actin cortex through distinct mechanisms, both can alter membrane mechanical state: hypotonic swelling increases membrane stretch, whereas actin depolymerization disrupts membrane–cortex coupling and membrane reservoir regulation. Notably, RhoA activation increased membrane-associated LifeAct-GFP signal (Fig. 1l), consistent with enhanced cortical actin assembly. Together with the corresponding changes in Flipper-TR lifetime across these conditions, these findings support a potential physical link between cortical cytoskeletal remodeling, membrane mechanics, and OMD organization.

### Rho activation modulates HCN channel function as a readout of OMD reorganization

Hyperpolarization-activated cyclic nucleotide-modulated (HCN) channels play an essential role in supporting autonomous neuronal firing^5,6^. The gating of HCN channels is sensitive to lipid compartmentalization, which makes them a functional reporter for detecting alterations in OMD^5,74^. Disruption of OMDs, such as by cholesterol depletion using β-cyclodextrin (β-CD), has been shown to markedly facilitate HCN channel gating, leading to faster activation and increased channel opening^5^. These changes in HCN gating contribute to enhanced neuronal excitability and are characteristic of sensory neurons under chronic neuropathic pain conditions^5,73^. In this context, we tested the role of RhoA signaling in modulating HCN channel function via OMDs. Using whole-cell patch clamp recordings, we measured native HCN currents in small-diameter DRG neurons from SNI rats by applying a series of hyperpolarizing voltage steps. To evaluate the role of RhoA signaling, we pharmacologically activated RhoA using CN03 and compared properties of HCN currents from treated and untreated controls (Fig. 7a). Ipsilateral nociceptor SNI neurons are known to exhibit disrupted OMDs and facilitated HCN channel opening, which contribute to their increased spontaneous activity^5^. We found that treatment with the Rho activator (1 μg/mL, >4 hours) partially reversed this pathological phenotype. HCN channel activation kinetics were markedly slowed, with the primary activation time constant (τ_1_) increasing from 75 ± 3 ms to 96 ± 6 ms (Fig. 7b). Although the half-maximal voltage (V_1/2_) for channel activation derived from the conductance-voltage (G-V) curve did not change (Fig. 7c), the slope factor of the G-V curve was significantly reduced, consistent with a restoration of OMDs in nociceptor DRG neurons (Fig. 7d). The reduced slope factor suggests increased total gating charge movement per channel during activation, consistent with membrane thickening caused by OMD expansion^5^. These findings demonstrate that RhoA activation supports OMD integrity, inhibits HCN channel function, and attenuates neuropathic hyperexcitability.

**Fig. 7.**
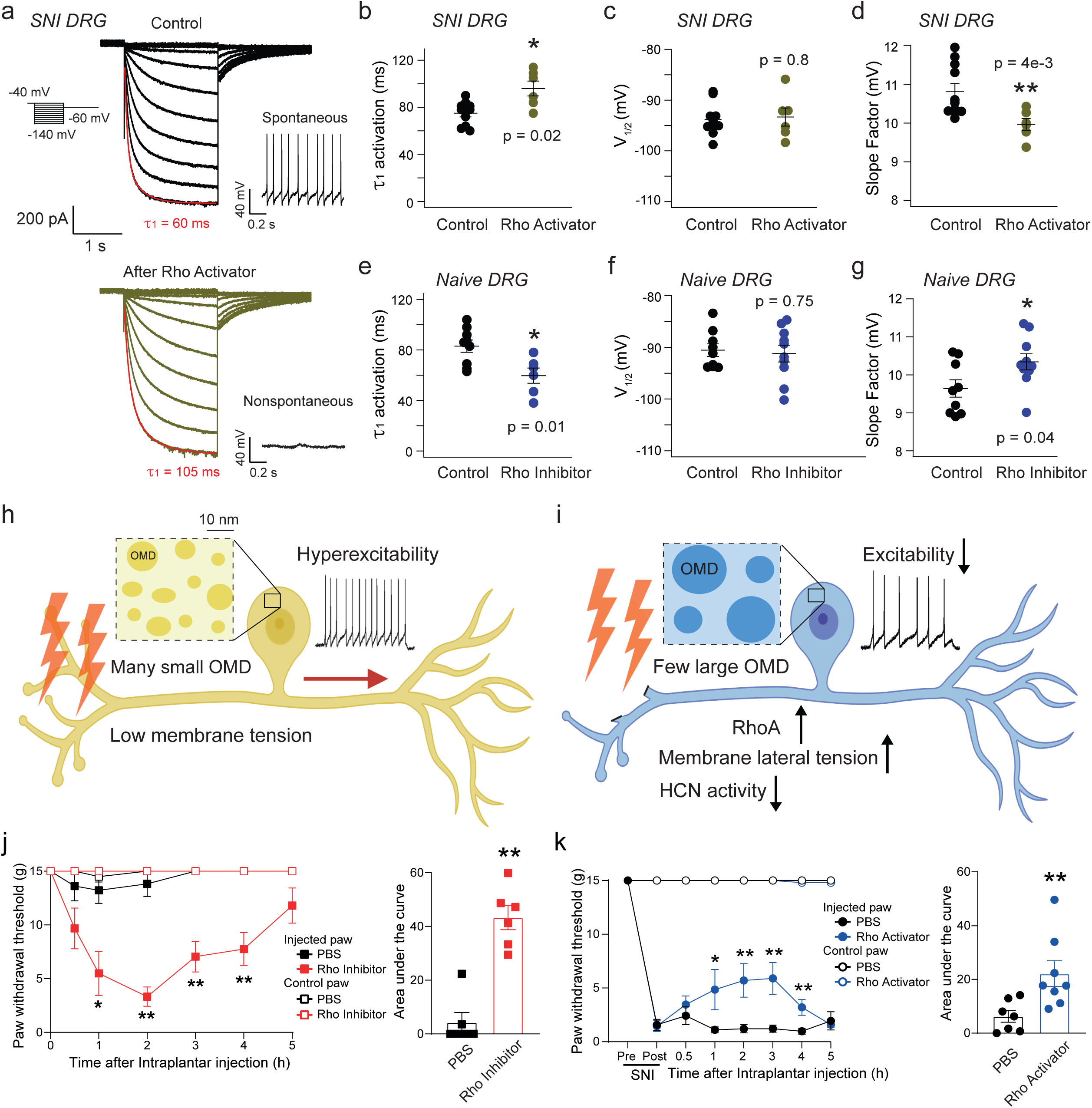
RhoA activation regulates HCN channel gating and protects against mechanical hypersensitivity in neuropathic pain. **a**, Representative HCN current traces showing that RhoA activation slows channel activation in SNI small DRG neurons. Red lines indicate double-exponential fits at −140 mV. Spontaneous action potential recordings from the same neuron are also shown. **b**-**d**, Summary of the parameters of Boltzmann fits to the G-V relationship for HCN channel activation before and after the RhoA activator treatment of SNI small DRG neurons: activation time constant (τ_1_) in b, V_1/2_ in c and slope factor (V_s_) in d, n = 10 patches, control and n = 6 patches, Rho activator, *p < 0.05, **p < 0.01, two-sided t-test. **e**-**g**, Summary of HCN channel activation before and after the RhoA inhibitor treatment of naïve small DRG neurons: activation time constant (τ_1_) in e, V_1/2_ in f and slope factor (V_s_) in g, n = 9 patches for the control, n = 6 for τ_1_ of HCN activation under the RhoA activator condition, and n = 9 (control) and 10 (Rho inhibitor) for V_1/2_ and V_s_ of the G-V curve fits for HCN activation, *p < 0.05, **p < 0.01, two-sided t-test. **h**-**i**, Proposed model illustrating how RhoA signaling suppresses neuronal hyperexcitability by increasing membrane mechanical state, promoting OMD remodeling, and reducing HCN channel activity. Under low membrane tension (neuropathic pain state, h), OMDs remain dispersed; increased membrane mechanical state following RhoA activation (i) drives OMD expansion and decreases HCN activity. **j**, Paw withdrawal thresholds in naïve male rats before and after intraplantar injection of a RhoA inhibitor (0.25 μg/50 μl) or PBS vehicle; the uninjected paw is shown for reference. *p = 0.02, 2e-3, 2e-3, 2e-3 at 1, 2, 3, 4 hr time points. The integrated area under the curve (AUC, 0–5 h) is shown in the adjacent summary graph. **p = 2e-3 versus vehicle, Mann–Whitney tests; n = 6 per group. **k**, Paw withdrawal thresholds in SNI rats before and after intraplantar injection of a RhoA activator (0.25 μg/50 μl) or PBS into the ipsilateral paw; the contralateral paw is shown for reference. *p = 0.02, 3e-3, 3e-3, 3e-3 at 1, 2, 3, 4 hr time points. The integrated AUC (0–5 h) is also shown. **p = 3.7e-3 versus vehicle, Mann–Whitney tests; n = 7 (PBS) and 8 (Rho activator), *p < 0.05, **p < 0.01.

To further support the OMD-mediated effect on HCN channels by RhoA, we investigated whether inhibition of RhoA might alter the channel gating in the opposite direction for naïve small DRG neurons. As expected, we found that inhibiting RhoA with C3 transferase (1 μg/mL for ∼4 hours) significantly accelerated HCN channel activation and increased the slope factor associated with the G-V activation curve (Fig. 7 e-g), recapitulating the β-CD-induced phenomenon of OMD disruption. Together, these results demonstrate that RhoA signaling critically stabilizes OMDs to regulate HCN gating and modulate action potential firing patterns.

### A proposed model linking RhoA activation to reduced neuronal hyperexcitability

We propose that, in neuropathic pain neurons, altered membrane mechanical state is associated with OMD disruption and hyperexcitability (Fig. 7h). RhoA and its downstream effectors, mDia and ROCK, function as a dynamic signaling network that modulates membrane mechanics and OMD organization. Through actin cytoskeletal remodeling, RhoA signaling may restore membrane mechanics and OMD organization (Fig. 7i), establishing a feedback mechanism that regulates ion-channel activity and neuronal excitability (Fig. 7i). In this model, HCN and other ion channels may act in concert to translate the restored membrane state into reduced excitability (Fig. 7i). Consistent with these cellular effects, intraplantar injection of the RhoA inhibitor, C3 transferase induced mechanical hypersensitivity in rats (Fig. 7j). Conversely, intraplantar injection of the RhoA activator, CN03 attenuated mechanical hypersensitivity in the SNI pain model (Fig. 7k), in agreement with the in vitro findings.

## Discussion

We propose that OMDs contribute to the spatial organization of membrane proteins and thereby influence their interactions and activity^1^. Changes in membrane order and nanoscale lipid compartmentalization have been linked to Alzheimer’s disease^11,12^, autoimmune disorders^1,14^, and neuropathic pain^5,73^. We previously showed that OMDs regulate the excitability of nociceptive DRG neurons, with smaller OMDs correlating with membrane hyperexcitability and increased action potential firing. In this study, we uncover a previously unknown link between the GTPase RhoA and neuropathic pain by examining how RhoA modulates OMDs. Using FLIM-FRET with OMD-associated probes, we achieved nanoscale sensitivity to monitor relative changes in OMD dimension following RhoA activation, accompanied by restoration of membrane mechanical state, supporting a role for RhoA in regulating membrane nanodomain organization.

We propose that during nerve injury and axon regeneration, RhoA activation serves as a protective mechanism against neuronal hyperexcitability by modulating OMDs. Previously, RhoA activation was reported to influence membrane excitability by regulating the trafficking of voltage-gated ion channels^75–77^. RhoA activation was also suggested to interact with IP_3_ receptors to modulate calcium signaling and the function of endothelial cells^78^. Nevertheless, the impact of RhoA activation specifically on nociceptor DRG neurons via ion channels has not been well studied. Our findings suggest that, beyond RhoA’s established role in ion channel trafficking, direct RhoA/OMD-mediated attenuation of HCN channel gating may further contribute to decreased excitability in the soma of DRG neurons upon RhoA activation. Furthermore, RhoA activation following nerve injury likely halts axon regeneration and restricts damage to the injured region to conserve energy for soma survival^21,79^, while simultaneously reducing somatic firing and alleviating the pain phenotype.

Our results support that RhoA activation promotes OMD reorganization through cytoskeletal remodeling and post-translational protein palmitoylation as underlying mechanisms. RhoA, a key regulator of actin cytoskeleton dynamics, modulates actomyosin contractility and cortical tension^17,20^. In neuropathic neurons, altered membrane mechanics—likely in part resulting from cholesterol loss^73^—may feedback to activate RhoA^80^, driving the coalescence of membrane domains into larger, more stable OMDs while concurrently restoring membrane mechanics. Our results show that acute RhoA activation was accompanied by rapid cortical actin remodeling mediated by mDia formin. In parallel, ROCK inhibition substantially reduced RhoA-dependent changes in OMD-associated FLIM-FRET signals in both tsA-201 cells and nociceptor DRG neurons, indicating that ROCK is a downstream effector of this pathway. Together, these findings support a model in which RhoA regulates the OMD organization through two major effectors mDia and ROCK, which together shape nanoscale lipid compartmentalization and influence neuronal excitability.

We postulate that the RhoA-promoted OMD remodeling may be dependent on palmitoylated proteins that preferentially localize to OMDs, such as GPI-anchored proteins, CD44^81,82^, caveolin^83^, and the PI(4,5)P_2_-binding protein GAP-43^84^. S-palmitoylation facilitates the localization of membrane proteins to OMDs^61,63^, and RhoA may reorganize the OMD localization of palmitoylated proteins by remodeling the cytoskeleton^62,85,86^. Additionally, RhoA activity likely regulates the subcellular localization or activity of palmitoyl acyltransferases that contain DHHC domains, thereby influencing the lipidation status of OMD-residing proteins^62,87^. Moreover, our findings support the hypothesis that RhoA plays a critical role in the clathrin/dynamin-independent, OMD-dependent form of endocytosis first described by Hilgemann and colleagues^28–30^. Notably, the actin cytoskeleton appears dispensable in the late stages of OMDE, suggesting that lipid compartmentalization and membrane tension dominate vesicle formation^30^. Altogether, these findings imply that RhoA acts as a molecular bridge between cytoskeletal forces, membrane mechanics, and lipid domain dynamics, coordinating mechanical and biochemical cues that promote membrane domain formation and endocytosis events.

Our results support a model where membrane mechanical environments help shape plasma membrane domain organization. Pathways that potentially elevate membrane tension, such as RhoA-dependent regulation of cortical actomyosin^69^, appear to promote nanoscale remodeling of OMDs. One possible interpretation is that, in living cells, increased lateral tension favors greater spatial clustering of OMDs, to reduce the cost of line-tension energy associated with domain boundaries^88,89^. Furthermore, the morphology of OMDs, which could adopt flat or dimpled configurations under different membrane tension states, was proposed to regulate their dimension and lateral organization^90^. Consistent with these ideas, acute hypotonic swelling or disruption of cortical F-actin, both of which alter membrane mechanics, produced changes in fluorescence-based measurements of membrane organization similar to those observed following RhoA activation. More broadly, these findings suggest that signaling pathways regulating membrane mechanical state can dynamically remodel membrane heterogeneity and organization, with potential consequences for membrane protein compartmentalization and neuronal excitability.

## Methods

### Molecular Reagents, Cell Culture, and Transfection

Frozen aliquots of tsA-201 cells (RRID: CVCL_2737), a type of human embryonic kidney (HEK) cell commonly used for studying membrane ion channels and electrophysiology, were acquired from Sigma-Aldrich (St. Louis, MO) and stored in liquid nitrogen. TsA-201 cells share key features of plasma membrane organization with neurons, including cholesterol-dependent membrane heterogeneity, allowing comparison of membrane regulatory mechanisms across cell types^5,6^. Together with the findings in DRG neurons, these results suggest that the observed effects may extend across multiple mammalian cell types.

Cells were cultured in Dulbecco’s modified Eagle’s medium (DMEM; Gibco), supplemented with 10% fetal bovine serum (FBS; Gibco) and 1% penicillin/streptomycin (Gibco), then stored in a humidified incubator at 37 degrees Celsius with 5% CO_2_. Transient transfection was performed on 50-90% confluent tsA-201 cells with the lipofectamine 3000 kit (Invitrogen, Carlsbad, CA, #L30000080). Mycoplasma testing was performed using the MycoFluor™ Mycoplasma Detection Kit (Invitrogen) and tested negative. The iLID system included LARG-mCherry-sspB (Addgene: #90459) and iLID-CaaX-ssrA (Addgene: #85680). DNA concentration was measured using a Nanodrop OneC spectrophotometer (Life Technologies, Grand Island, NY). The cholera toxin subunit B-Alexa Fluor^TM^ 488 and 647 conjugates were purchased from ThermoFisher Scientific and administered to cultured cells at a concentration of 20 nM for 5-10 minutes^5^. L10-CFP, L10-YFP, S15-CFP, and S15-YFP are gifts from Prof. Bertil Hille (University of Washington, Seattle). The membrane-permeable Rho activator II and inhibitor I were all purchased from Cytoskeleton, Inc (Denver, CO). The Flipper-TR biosensor of membrane mechanics (#SC020) was from Spirochrome (University of Geneva, Switzerland) and applied at 1 μM for ∼40 mins before imaging. Latrunculin A was purchased from ThermoFisher (#L12370). 2-bromopalmitate (#21604) was purchased from Sigma (St. Louis, MO) and was prepared at 5:1 ratio mixed with fatty-acid free bovine serum albumin (#A8806, Sigma). BSA-palmitate was from Cayman Chemical (#29558; 6:1 fatty acid to BSA ratio). ROCK inhibitor Y-27632 (#Y0503) was from Sigma.

### Fluorescence Lifetime Measurement and confocal FLIM-FRET

Digital frequency-domain fluorescence lifetime imaging^38,91^ was performed with a Q2 laser scanning confocal system equipped with a FastFLIM data acquisition module (ISS, Inc, Champaign, IL), and two hybrid PMT detectors (Model R10467-40, Hamamatsu USA, Bridgewater, NJ) at ambient room temperature. An Olympus IX73 inverted microscope (Olympus America, Waltham, MA) is coupled to the confocal system. Both the frequency domain (instantaneous distortion-free phasor plotting) and time-domain decay information are included in the fluorescence lifetime measurements. Simultaneous two-color imaging was performed with dichroic cubes, one equipped with a 50/50 beam splitter and the other with various long-pass filters. A supercontinuum laser with a wavelength range of 410 to 900 nm, a repetition rate of 20 MHz, and a pulse duration of 6 ps was used for fluorescence excitation. Specifically, AF-488 was excited at 488 nm and mCherry was excited at 561 nm. To detect the emission for AF-488 (or GFP) and mCherry (or mRuby), an emission cube filter consisting of a 525/40 nm filter for AF-488, a 552 nm long-pass dichroic mirror, and a 593/40 nm filter for mCherry was employed. For L10 FRET measurements, CFP-and YFP-tagged probes were co-transfected to the same cells. Cells expressing both fluorophores were randomly selected for imaging, whereas cells displaying substantial imbalance in CFP and YFP fluorescence were excluded based on visual inspection^5,92^. To detect the emission for CFP and YFP, an emission filter cube with a 475/28 nm filter for CFP, a 495 nm long-pass dichroic mirror, and a 542/27 nm filter for YFP was used. Only the donor lifetime was measured for phasor FLIM-FRET either in the presence or absence of the FRET acceptor. Optogenetic activation was performed with a supercontinuum laser using 445 nm or 488 nm excitation. The lifetime imaging of Flipper-TR was using 488 nm excitation and emission was collected with the 593/40 nm filter.

Confocal images of 256 x 256 pixel frame were obtained under each experimental condition, with a 100 μm motorized variable pinhole and a pixel dwell time of 0.1-0.4 ms. Acquired pixels were examined using the phasor plot. The VistaVision software (https://iss.com/software/vistavision) was used for image processing, display, and acquisition and allows for the specification of parameters such as pixel dwell time, image size, and resolution. The fitting algorithm and multi-phasor analysis were used for FLIM analysis. A median/gaussian smoothing filter and intensity-threshold filter were applied to enhance the display and analysis of membrane-localized lifetime species. The smoothing filter was applied uniformly across all experiments to enhance the precision of phasor-based lifetime localization. For all datasets, membrane regions of interest (ROIs) were defined based on the corresponding intensity images, using a consistent ROI selection strategy across all experimental conditions. Importantly, all membrane-localized lifetime phasors were verified to be stable over a range of reasonable threshold values, indicating that reported differences were not dependent on minor variations in pixel selection. Phasor FLIM can differentiate between signals originating from the background outside of the membrane and those within the membrane. For CTxB experiments, internalized signal was typically minimal under our experimental conditions at room temperature (Supplementary Fig. 1) but, when present, was distinguishable as intracellular puncta with shorter fluorescence lifetime values^5^. These pixels typically form a separate cluster in phasor space, consistent with our previous characterization^5^, and were excluded from membrane ROI selection. This approach does not directly measure absolute OMD dimension but instead provides a sensitive measure of relative changes in domain dimension, enabling quantitative comparisons across experimental conditions.

The VistaVision software was used for determination of phase delays (φ) and modulation ratios (m) of the fluorescence signal in response to an oscillatory stimulus with a frequency of ω. Sine (S) and cosine (S) Fourier transforms of the phase histogram were performed with first harmonic and displayed at the 40-MHz modulation frequency, with consideration for the instrument response function (IRF) calibrated with Atto 425 (lifetime: 3.6 ns) and rhodamine 110 (lifetime: 4 ns) measured in solution, for 445 nm and 488 nm excitations, respectively. The FLIM imaging of Flipper-TR was calibrated using rhodamine B (lifetime: 1.68 ns). The equations used were standard for frequency domain FLIM^40,41^, including 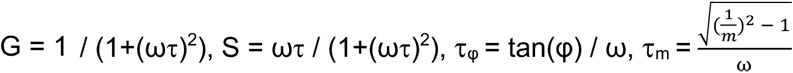.

The FRET trajectory function in the VistaVision software allowed for analysis of FLIM-FRET measurements^37^. Three parameters were adjusted to ensure accurate results and optimize the fitting trajectory through both the unquenched donor (donor alone) and donor species affected by FRET. These parameters include the background contribution in the donor sample, unquenched donor contribution in the FRET sample, and the background contribution in the FRET sample. Background levels for both the donor sample and FRET sample were typically under 4%, which corresponds with the very low emission intensity observed in untransfected cells.

The RhoA activity was monitored using the genetically encoded FluoSTEP-RhoA biosensor^48^ (Addgene: #181860 and #181859). The biosensor consists of two components: full-length RhoA fused to the GFP11 peptide and a reporter module containing the complementary GFP1–10 fragment of superfolder GFP (sfGFP), a circularly permuted RhoA-binding domain from Protein Kinase C-Related Kinase 1 (cpPKN), and mRuby2. GFP1–10 and GFP11 correspond to β-strands 1–10 and β-strand 11 of sfGFP, respectively, and they reconstitute a functional FRET donor upon co-expression^48^. Activated GTP-bound RhoA binds the cpPKN domain, resulting in a change in FRET between reconstituted sfGFP and mRuby2^48^. Cells were co-transfected with both biosensor components and imaged by live-cell FLIM. FRET efficiency was quantified using the same FLIM-FRET approach described above. The alternative FRET-based RhoA biosensor pTriExRhoA2G (Addgene: #40176) exhibits opposite directional change of FRET during RhoA activation, compared to FluoSTEP-RhoA (Supplementary Fig X). The FRET of biosensor pTriExRhoA2G increases with RhoA activation^49^. The localization-based biosensor of RhoA mNeonGreen-3xrGBD^59^ was obtained from Addgene (#176091) and is compatible with the iLID system.

Cholesterol level at the plasma membrane was assessed using the genetically encoded cholesterol sensor eGFP-GRAM-W^6,93^. The tsA cells expressing GRAM-W were imaged by fluorescence polarization microscopy, and fluorescence emission was collected in parallel (I_∥_) and perpendicular (I_⊥_) polarization channels following excitation with linearly polarized light. The fluorescence anisotropy (r) at the plasma membrane was calculated using the equation: r = (I_∥_ − I_⊥_) / (∥ + 2*I_⊥_)^6^. The lack of change in anisotropy suggests that homo-FRET between the cholesterol sensors remained largely unchanged, indicating no significant alteration in the relative level of accessible cholesterol at the plasma membrane^6^.

Three buffer solutions were formulated to achieve specific osmolarity for Flipper experiments. A hypotonic buffer (154 mmol/kg) was prepared by dissolving 37 mM NaCl, 5 mM KCl, 1 mM MgCl_2_, 2 mM CaCl_2_, 10 mM D-(+)-Glucose, and 20 mM HEPES. An isotonic buffer (326 mmol/kg) was prepared similarly but with 137 mM NaCl. A hypertonic buffer (493 mmol/kg) was then made by adding 150 mM sucrose to the isotonic buffer composition. The pH of all solutions was adjusted to 7.4, and they were subsequently sterile-filtered and stored at 4°C.

### Animals and Dorsal Root Ganglion (DRG) Neuron Preparation

All procedures involving animals were approved by the Institutional Animal Care and Use Committee (IACUC) of Saint Louis University and adhered to the National Institutes of Health guidelines. Adult pathogen-free Sprague-Dawley rats (100 – 250 g, Envigo, Placentia, CA) were housed under a 12-hour light/dark cycle (lights on at 06:00 AM) in a temperature-controlled room (23 ± 3°C) with ad libitum access to food and water. All behavioral experiments were conducted by investigators blinded to treatment conditions.

Primary DRG neurons were isolated from naïve (100 g) or adult (200 – 250 g) rats subjected to nerve injury. Animals were deeply anesthetized, and the dorsal skin and underlying musculature were carefully dissected to expose the vertebral column. Dorsal root ganglia (DRGs) were isolated by removing the vertebral bone overlying the spinal cord, followed by excision and trimming of the DRG roots. Isolated DRGs were enzymatically digested in 3 mL of sterile, bicarbonate-free, serum-free DMEM (Catalog# 11965, Thermo Fisher Scientific, Waltham, MA) containing 3.125 mg/mL neutral protease (Catalog# LS02104, Worthington Biochemical Corp., Lakewood, NJ) and 5 mg/mL collagenase Type I (Catalog# LS004194, Worthington). Tissue digestion was carried out at 37°C for 45 minutes with gentle agitation. Following digestion, cells were centrifuged and resuspended in DRG culture medium consisting of DMEM supplemented with 10% fetal bovine serum, 30 ng/mL nerve growth factor, and 1% penicillin/streptomycin. Cells (∼1.5 × 10^6^) were plated onto 12-mm coverslips pre-coated with poly-D-lysine and laminin and maintained at 37°C in a humidified incubator with 5% CO_2_ until experimentation.

### Spared Nerve Injury (SNI) Surgery and Behavioral Testing

Rats were anesthetized using 5% isoflurane for induction and maintained at 2.5% during surgery. Animals were placed on a heating pad to maintain core body temperature, and areflexia was confirmed prior to incision. The surgical site on the right hindlimb was cleaned with chlorhexidine (applied twice), and a 2–3 cm incision was made along the lateral thigh. The biceps femoris muscle was bluntly retracted to expose the sciatic nerve and its three terminal branches. The common peroneal and tibial nerves were tightly ligated using 4–0 silk sutures and transected, while the sural nerve was left intact. The muscle was closed with 5–0 absorbable suture, and the skin was closed with surgical clips. Topical antibiotic ointment was applied, and animals were allowed to recover for 10 days before behavioral assessments.

Von Frey mechanical sensitivity testing was used to assess tactile allodynia. Testing was conducted in a blinded manner in a quiet, temperature-controlled room. Rats were acclimated to the testing environment in individual Plexiglas chambers placed on a wire mesh platform. Calibrated von Frey filaments were applied to the lateral plantar surface of the hindpaw, and the 50% paw withdrawal threshold was determined using the up-down method.

### Patch-Clamp Electrophysiology

Electrophysiological recordings were performed in whole-cell configuration using an EPC10 amplifier and PATCHMASTER software (HEKA Elektronik), with data acquired at a sampling rate of 5 kHz. Recording pipettes were pulled from borosilicate glass capillaries using a P1000 puller (Sutter Instruments, Novato, CA) and had a resistance of 2.5-4 MΩ when filled. Precise positioning of pipettes was achieved using a Sutter MP-225A motorized micromanipulator. All recordings were conducted at ambient room temperature.

To record HCN channel currents in dorsal root ganglion (DRG) neurons, the internal pipette solution contained (in mM): 10 NaCl, 137 KCl, 4 Mg-ATP, 10 HEPES, and 1 EGTA, pH adjusted to 7.3 with KOH. The inclusion of 4 mM Mg-ATP was sufficient to preserve intracellular phosphoinositide levels, particularly PI(4,5)P₂, throughout the duration of recordings (52, 53). The extracellular solution consisted of (in mM): 154 NaCl, 5.6 KCl, 1 MgCl_2_, 1 CaCl_2_, 8 HEPES, and 10 D-glucose, titrated to pH 7.4 with NaOH. Series resistance was routinely maintained below 10 MΩ and compensated by 40%–70%. For the action potential measurements, whole-cell current clamp mode of PATCHMASTER was used instead of the voltage-clamp configuration for HCN current measurements. Spontaneous firing was measured with 0 pA current injection and defined as the occurrence of at least one action potential during the 1.4-s recording period. Input resistance was determined from the voltage responses to small hyperpolarizing current injections using Ohm’s law. For the current injection-evoked action potential firing, membrane potential was initially held at −60 mV. During current injection, neurons exhibited firing that adapted at higher amplitudes, likely due to sodium channel inactivation.

HCN current activation properties were assessed by measuring tail currents at −60 mV, following a series of test voltages from −40 mV to −140 mV in 10 mV increments. Leak-subtracted tail currents were normalized to the maximal amplitude and plotted to generate conductance-voltage (G–V) relationships. These data were fitted with a Boltzmann function: G/G_max_ = 1/(1 + exp[(V − V_1/2_)/V_S_]), where V is the test potential, V_1/2_ is the voltage at which half-maximal activation occurs, and V_S_ is the slope factor. The slope factor was defined as V_S_ = RT/zF, where R is the gas constant, T is the absolute temperature (297 K), F is the Faraday constant, and z is the effective gating charge.

### Statistics and Reproducibility

Data are presented as mean ± SEM. The number of observations (n) represents the number of cells or patches analyzed. The number of independent biological replicates (independent transfections, cell cultures, neuronal preparations, or animals) is reported in the Reporting Summary. Experiments were performed using cells obtained from at least 2–5 independent preparations, with both control and treatment conditions included in each preparation. Because of the limited number of biological replicates, formal normality testing was not used as the primary criterion for statistical analysis. Two-tailed Student’s t-tests were used for comparisons between two groups. One-way ANOVA with a Tukey’s post hoc tests was used for multiple-group comparisons. Paired t-tests were used for same-cell measurements before and after treatment. A value of *p < 0.05 or **p < 0.01 was considered statistically significant.

## Supporting information

This file includes Supplementary Figures 1-12, Supplementary Table 1

## Data Availability

Source data supporting Figs. 1 - 7 and supplementary figures are provided as a Source Data file in figshare.com as: https://doi.org/10.6084/m9.figshare.31350442, and additional data are in the Supplementary Information.

## Code Availability

This paper does not include any computational codes.

## Acknowledgements

We thank Erin N. Lessie and Lyuba Salih for technical support, and Kyle S. McCommis for sharing chemicals. This research is supported by National Institute of General Medical Sciences R35GM154778 grant (to G.D.), R56HL169176 (to G.D.), and by the Doisy Fund (to G.D. and L.J.H.) of the Edward A. Doisy Department of Biochemistry and Molecular Biology at Saint Louis University School of Medicine. The Moutal lab is supported by startup funds from the Department of Pharmacology and Physiology, Saint Louis University, Institute for Drug & Biotherapeutic Innovation seed grant, National Institute of Neurological Disorders and Stroke R01NS119263 grant. Cartoons in figures were created using BioRender.

## Author Contributions

G.D., A.M., and D.W.H. conceptualized and designed the research. S.S., L.J.H., M.D.J.N. and G.D. performed experiments and analysis for fluorescence imaging and electrophysiology. N.L.M., A.D.L., W.G., and G.D. performed pilot experiments. A.M., C.G., N.L.A.D., G.G., and H.A. performed DRG neuron preparations and rat studies. A.M. supervised rat studies. G.D. wrote the manuscript.

