## Supplementary material for "Dynamic RhoA-Mediated Remodeling of Ordered Membrane Domains": This file includes Supplementary Figures 1-12, Supplementary Table 1

**Supplementary information for "Dynamic RhoA-Mediated Remodeling of Ordered Membrane Domains"**

**Supplementary Fig. 1**

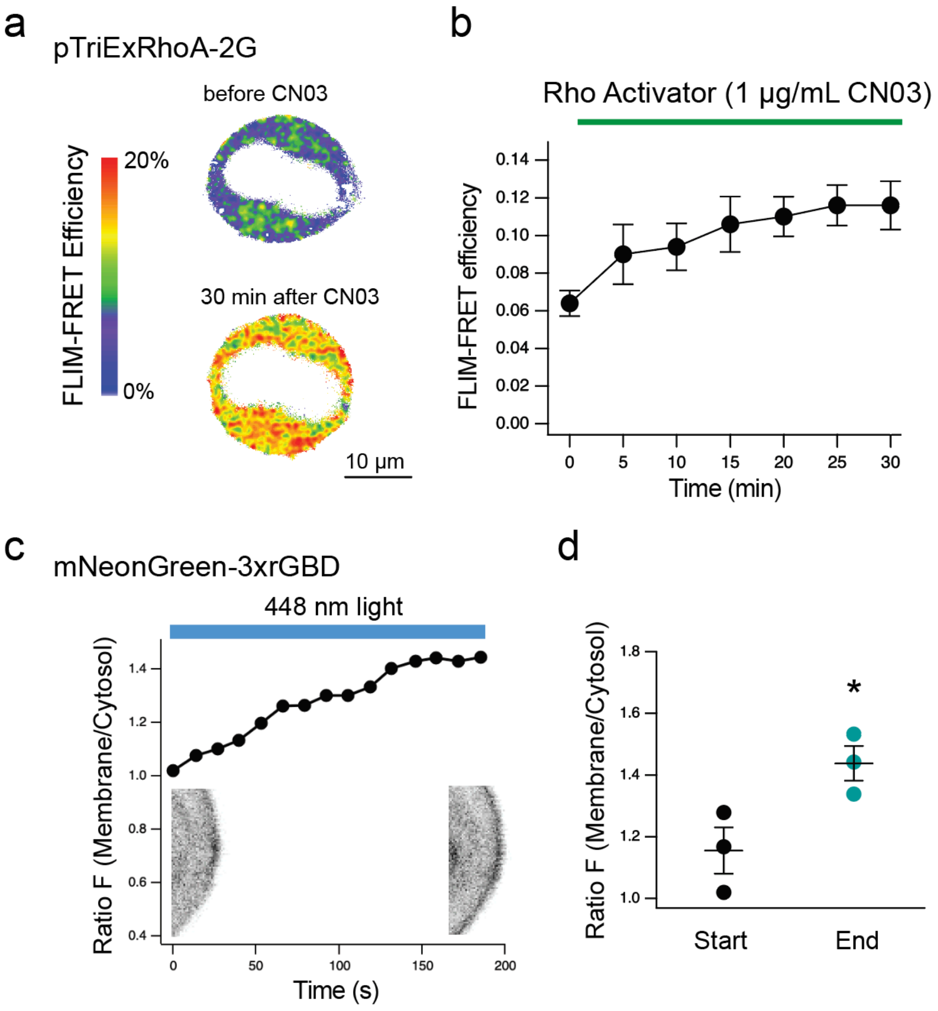


**Fig. S1. Alternative RhoA sensors report the pharmacological or iLID-mediated RhoA activation.**

**a-b**. The FRET-based RhoA sensor RhoA-2G showed an increase in FRET during CN03 application. Representative FRET heatmap images of tsA cells expressing the sensor are shown in a, and the corresponding summary time course of the FRET increase is shown in b. n = 5, **p = 0.01 at 30 min time point, mean ± s.e.m., paired t-test.

**c-d**. The translocation-based RhoA sensor mNeonGreen-3xrGBD was used to monitor RhoA activation during iLID-mediated light stimulation. Representative images of tsA cells at the beginning and end of 488-nm illumination, together with the time course of the membrane-to-cytosol fluorescence intensity ratio, demonstrate increased membrane localization of the sensor. The fluorescence intensity ratio before and after light stimulation is summarized in d. n = 3, mean ± s.e.m., p = 0.03, paired t-test**.**

**Supplementary Fig. 2**

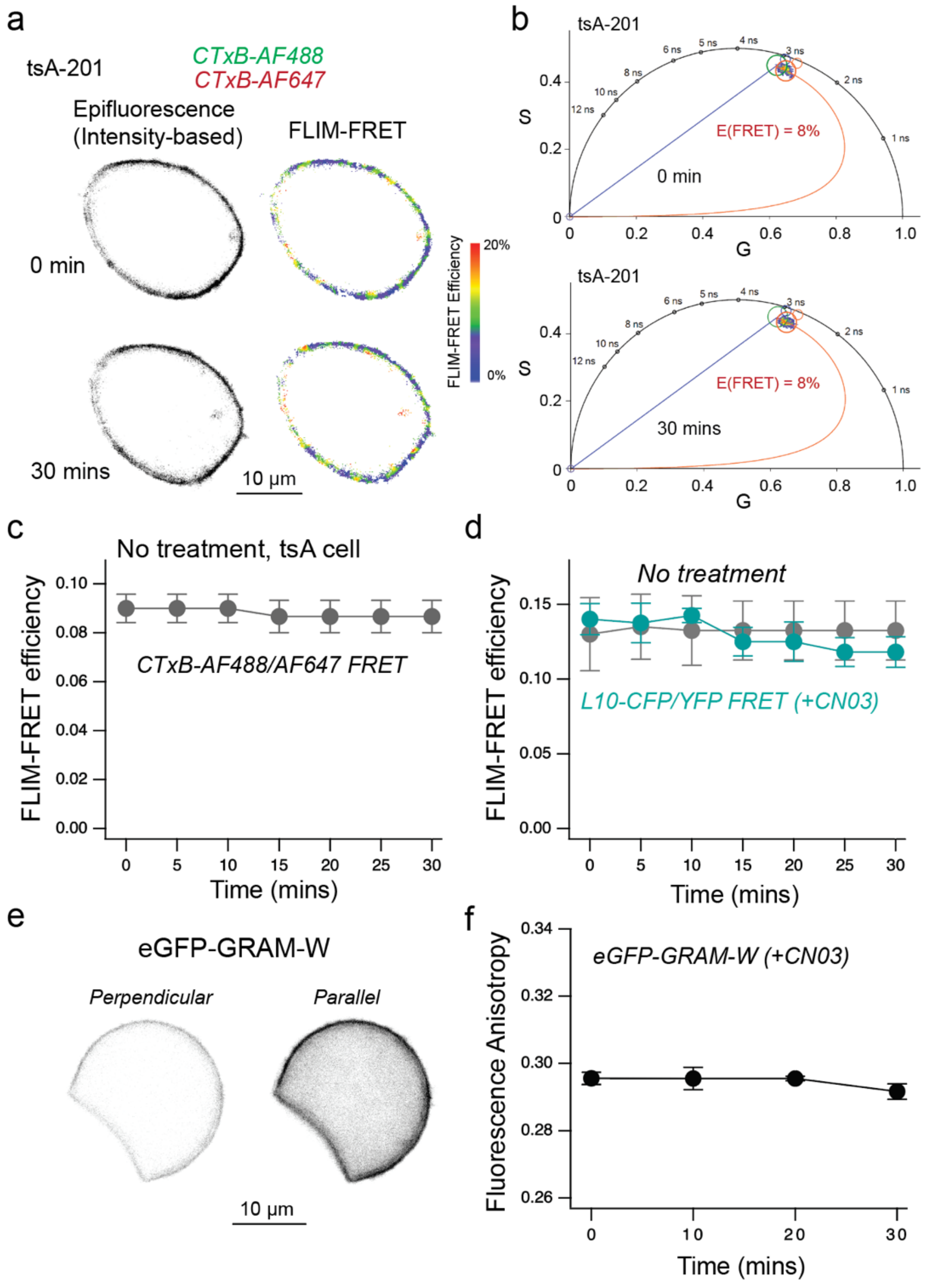


**Fig. S2. Control experiments for CTxB-based and L10-based FRET measurements during acute CN03 application.**

**a-b**. Representative intensity-based epifluorescence images and FLIM-FRET heatmaps (a), together with corresponding phasor plots (b), from untreated time-control cells at the start of imaging (0 min) and after 30 min at room temperature. No significant change in FRET efficiency was observed over the imaging period, and endocytosis of CTxB was minimal.

**c**. Time course of FRET efficiency in untreated control cells for the CTxB-based FRET assay ($n = 3$, $p = 0.42$ at 30 min, two-sided paired t-test).

**d**. Time course of FRET efficiency measured using the L10-based FRET pair in cells treated with CN03 (1 μg/mL) or in untreated time-control cells. FRET efficiency was significantly

different with p = 0.01 at 30 min for the CN03-treated cells (n = 5), whereas no significant change was observed in the time-control cells (p = 0.64, n = 4), two-sided paired t-test.
**e-f**. The cholesterol-sensitive probe GRAM-W showed no detectable change in fluorescence anisotropy during CN03 (1 μg/mL) application, suggesting that the membrane density of free cholesterol was not significantly altered. Representative perpendicular and parallel fluorescence emission intensities from a tsA cell are shown in e, and the corresponding time course of GFP anisotropy is summarized in f. No statistically significant change in anisotropy was observed over the course of CN03 application, p = 0.99 at 10 min, 0.97 at 20 min, and 0.2 at 30 min, two-sided paired t-test, mean ± s.e.m., n = 4.

**Supplementary Fig. 3**

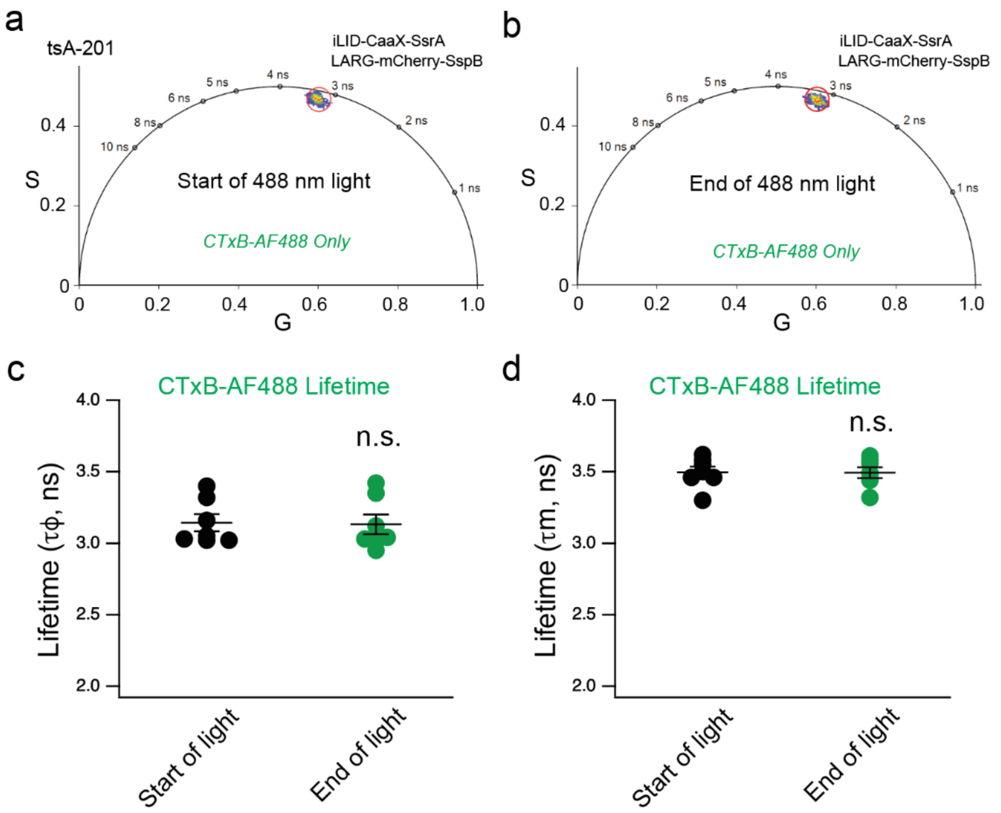


**Fig. S3. Optogenetic activation of RhoA does not alter CTxB-AF488 lifetime in the absence of FRET acceptor.**

**a-b**. Representative phasor plots of tsA-201 cells labeled with the CTxB-AF488 donor alone, shown before (a) and after (b) 488-nm light activation of the iLID optogenetic system. The cursor position (red) on the phasor plot indicates the donor species' phase lifetime ($\tau\varphi$) and modulation lifetime ($\tau m$).

**c-d**. Quantification of CTxB-AF488 phase lifetime (c) and modulation lifetime (d), comparing values at the start and end of 488 nm light exposure. Mean ± s.e.m., n = 7, two-sided t-test, n.s., no statistically significant difference.

**Supplementary Fig. 4**

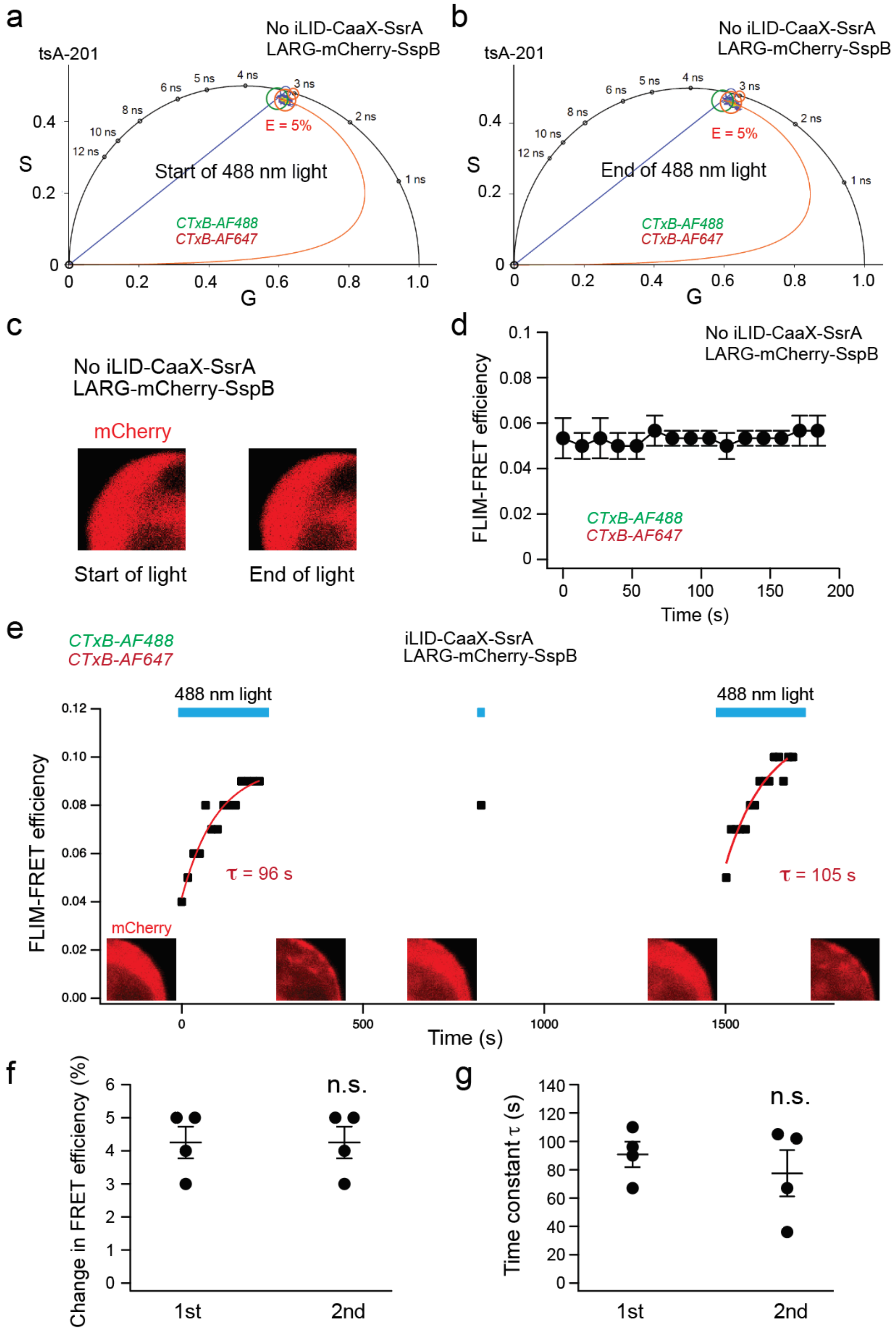


**Fig. S4. Control and reversible activation–deactivation experiments using optogenetic iLID system.**

**a-b**. Representative phasor plots of tsA-201 cells labeled with CTxB-AF488 and CTxB-AF647 before (a) and after (b) 488 nm illumination in cells lacking the iLID-CaaX membrane anchor. No change in CTxB-based FRET was observed.

**c**. Representative images of tsA-201 cells expressing LARG-mCherry-SspB alone (without cotransfection of iLID-CaaX), showing no membrane translocation of mCherry following 488 nm illumination.
**d**. Time course of CTxB-based FRET in cells lacking the iLID-CaaX membrane anchor, demonstrating no detectable FRET change upon 488 nm illumination (n = 3, p = 0.4 before vs. after).
**e**. An activation–deactivation paradigm demonstrating the reversible increase in CTxB-based FRET induced by repeated 488 nm illumination. Following the initial light pulse, CTxB-based FRET gradually recovered after the light was turned off, while a second 488 nm stimulation elicited a comparable increase in FRET. A brief 488 nm pulse applied during the recovery phase confirmed that the iLID system had already recovered, even though the CTxB-based FRET signal had not yet fully recovered. The corresponding mCherry fluorescence is shown to monitor the activation state of the iLID system throughout the experiment. The red curves indicate mono-exponential fits of the FRET increase.
**f-g**. Summary of the reversible iLID experiments shown in panel e, comparing the changes in FLIM-FRET efficiency (f) and the monoexponential time constants (g) elicited by the first and second rounds of 488 nm stimulation, mean ± s.e.m.; n = 4, two-sided paired t-test, n.s., no statistically significant difference.

**Supplementary Fig. 5**

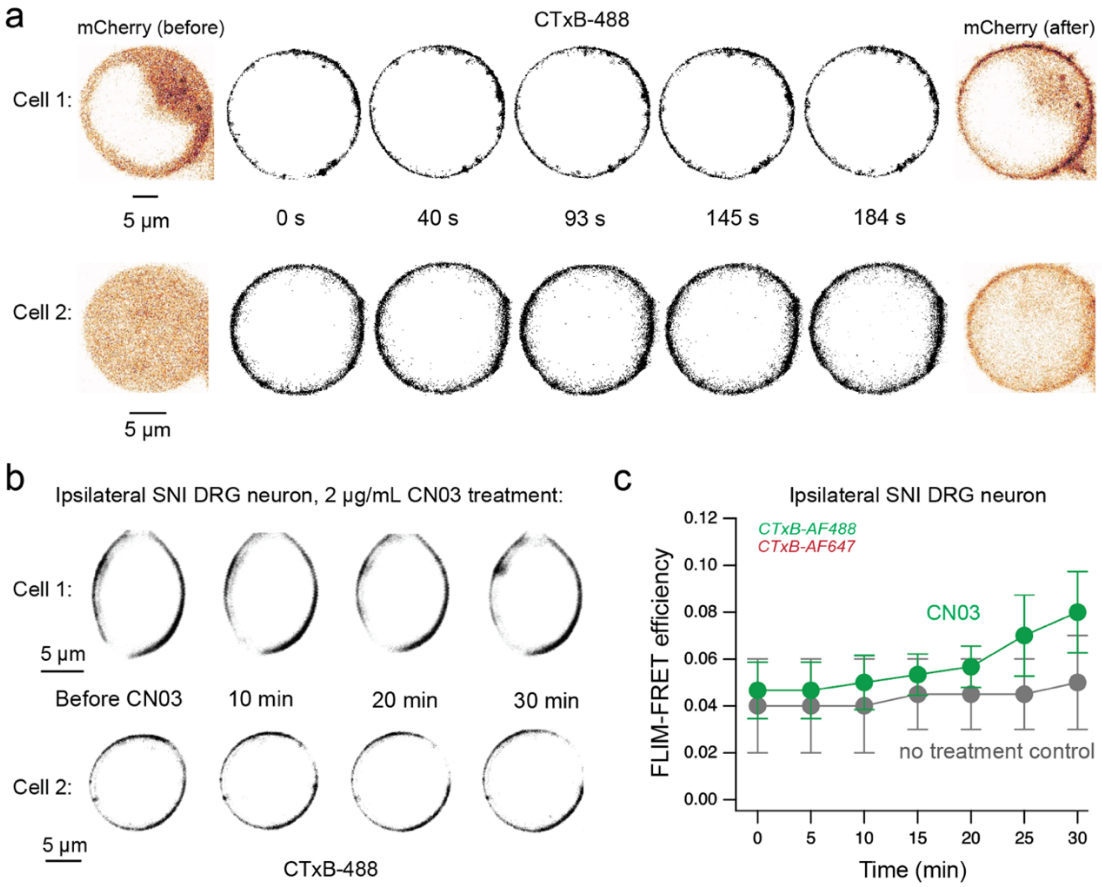


**Fig. S5. Morphological changes in tsA-201 cells and small DRG neurons during RhoA activation.**

**a**. Representative tsA-201 cells labeled with CTxB-AF488 showing plasma membrane morphology during optogenetic RhoA activation. mCherry fluorescence before and after 488-nm illumination was used to monitor the membrane translocation of mCherry-LARG.

**b**. Representative ipsilateral SNI DRG neurons labeled with CTxB-AF488 and CTxB-AF647 during CN03 treatment, illustrating changes in cell morphology following RhoA activation, as visualized by membrane-localized AF488 fluorescence.

**c**. Summary graph showing the increase in CTxB-based FRET following 30 min of treatment with 2 µg/mL CN03. In contrast, the untreated control exhibited minimal change over the same time period. $p = 0.038$ at 30 min ($n = 3$) after CN03 treatment, mean ± SEM, two-sided paired t-test.

**Supplementary Fig. 6**

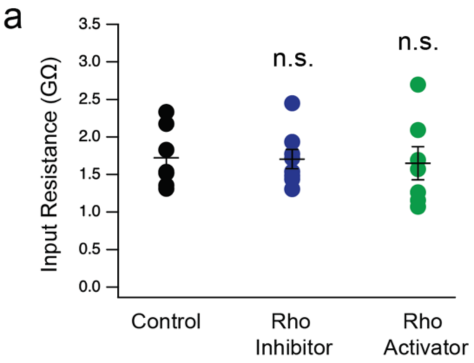


**Fig. S6. Input resistance of naïve DRG nociceptor neurons after Rho activation and inhibition.**

**a**. Overnight treatment with the Rho inhibitor C3 transferase (1 µg/mL) or the Rho activator CN03 (1 µg/mL) did not significantly alter the input resistance of naïve DRG neurons, n = 7 for the control and n = 8 for the Rho inhibitor, n = 7 for the Rho activator. Data shown as mean ± SEM, two-sided t-test.

**Supplementary Fig. 7**

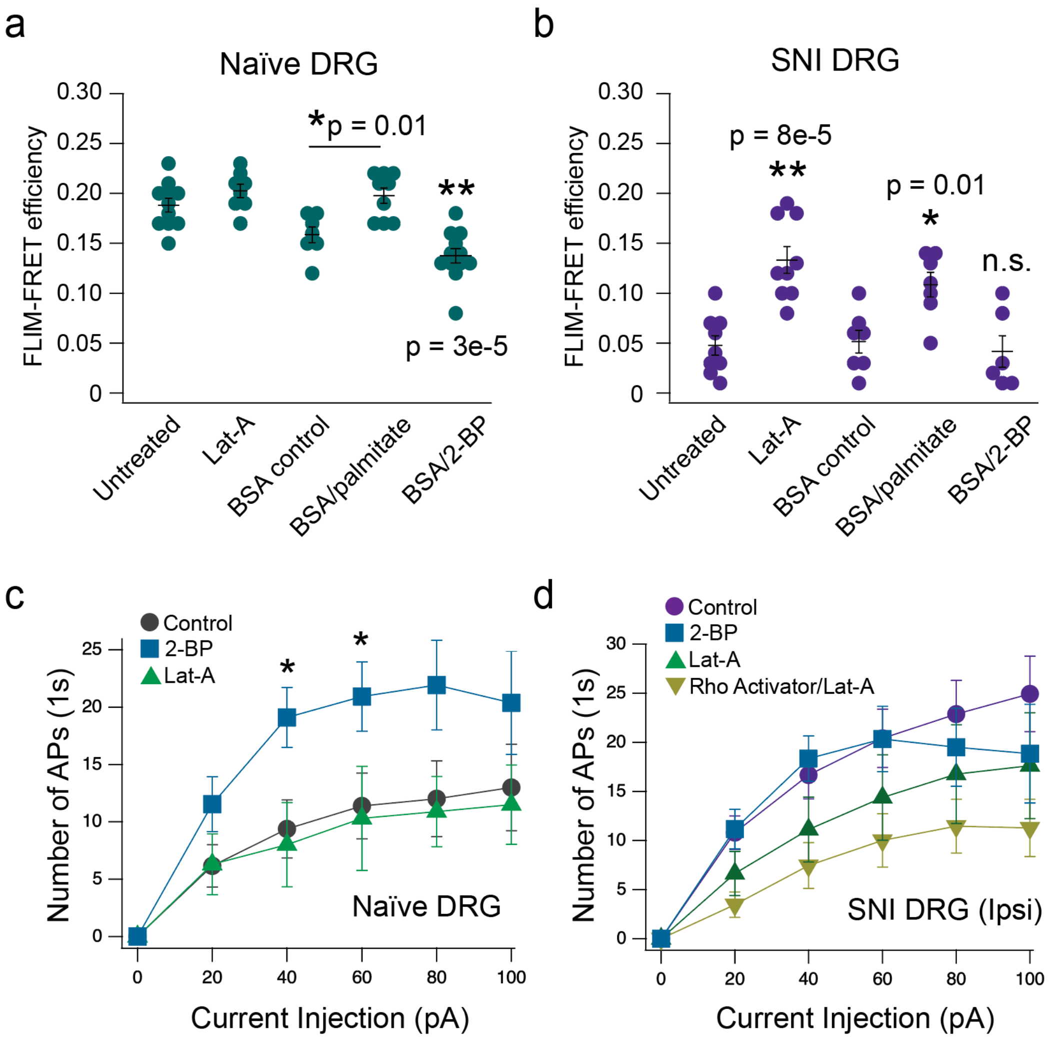


**Fig. S7. Actin cytoskeleton and protein palmitoylation regulate OMD organization and neuronal excitability in naïve and SNI nociceptor neurons.**

**a**. Baseline CTxB-based FLIM–FRET efficiencies of naïve nociceptor DRG neurons following treatment with Lat-A (n = 8), BSA (n = 7), BSA/palmitate (n = 9), or BSA/2-BP (n = 12), relative to the control (n = 11), one-way ANOVA. **b**, Corresponding CTxB-based FRET efficiencies for ipsilateral SNI neurons across the same pharmacological manipulations (control: n = 9; Lat-A: n = 9; BSA: n = 7; BSA/palmitate: n = 7; BSA/2-BP: n = 6). **c**, Summary of current injection-elicited AP firing in naïve DRG neurons after alteration of protein palmitoylation or actin cytoskeleton disruption (control: n = 13; 2-BP: n = 11; Lat-A: n = 10; mean ± s.e.m.; p = 0.01 at 40 pA and p = 0.03 at 60 pA, two-sided t-test between control and 2-BP conditions). **d**, Summary of action potential firing in ipsilateral SNI neurons elicited by current injection, following manipulation of protein palmitoylation, or comparing conditions with and without Rho activator treatment prior to actin cytoskeleton disruption (2-BP: n = 6; Lat-A: n = 8; Rho activator/Lat-A: n = 11); mean ± s.e.m.; no significance comparing Lat-A and Rho activator/Lat-A conditions, two-sided t-test.

**Supplementary Fig 8**

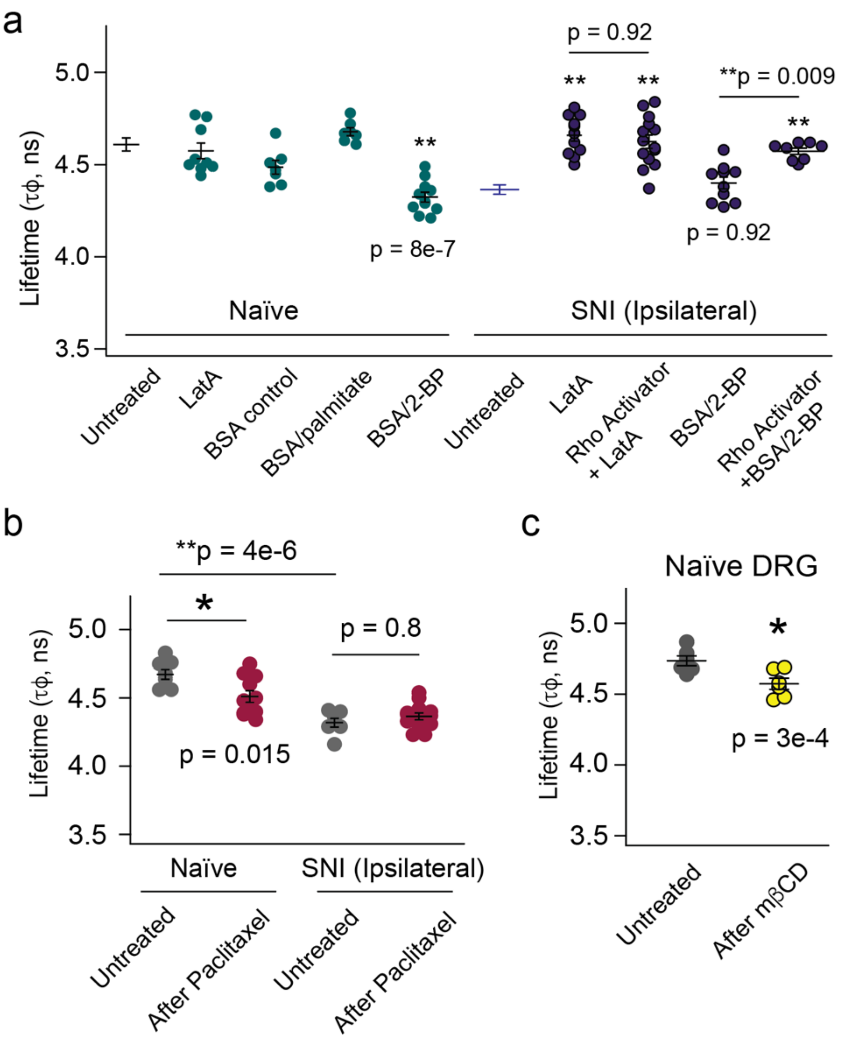


**Fig. S8. Actin cytoskeleton, palmitoylation, microtubule stabilization and lipid composition differentially regulate membrane mechanics in nociceptor DRG neurons.**

**a**. Flipper-TR lifetime measurements indicate that the membrane mechanical state is differentially regulated by actin cytoskeleton and protein palmitoylation, in parallel with their effects on the OMD size. In naïve DRG neurons, palmitoylation inhibition with 2-BP decreased Flipper lifetime, whereas actin disruption with Lat-A had little effect (n = 7-11). In ipsilateral SNI nociceptors, however, Lat-A increased Flipper lifetime, while 2-BP produced no additional reduction. Activation of RhoA increased Flipper lifetime in an actin-dependent manner, as this effect was eliminated by Lat-A but not by 2-BP, suggesting palmitoylation was acting upstream of RhoA signaling (n = 8-11). **b**. Stabilization of microtubules with paclitaxel (1 μM, overnight) reduced Flipper lifetime in naïve DRG neurons but had minimal effect in ipsilateral SNI neurons, suggesting regulation of the membrane mechanical environment distinct from actin-dependent mechanisms. Mean ± s.e.m., n = 7 for the control and n = 13 with paclitaxel for SNI; n = 8 for the control and n = 11 with paclitaxel for naïve neurons. **c**. Cholesterol depletion with MβCD decreased Flipper lifetime in naïve DRG neurons. Mean ± s.e.m., n = 5, one-way ANOVA for multiple conditions, and two-sided t-test for comparing two conditions. *p < 0.05, **p < 0.01.

**Supplementary Fig 9**

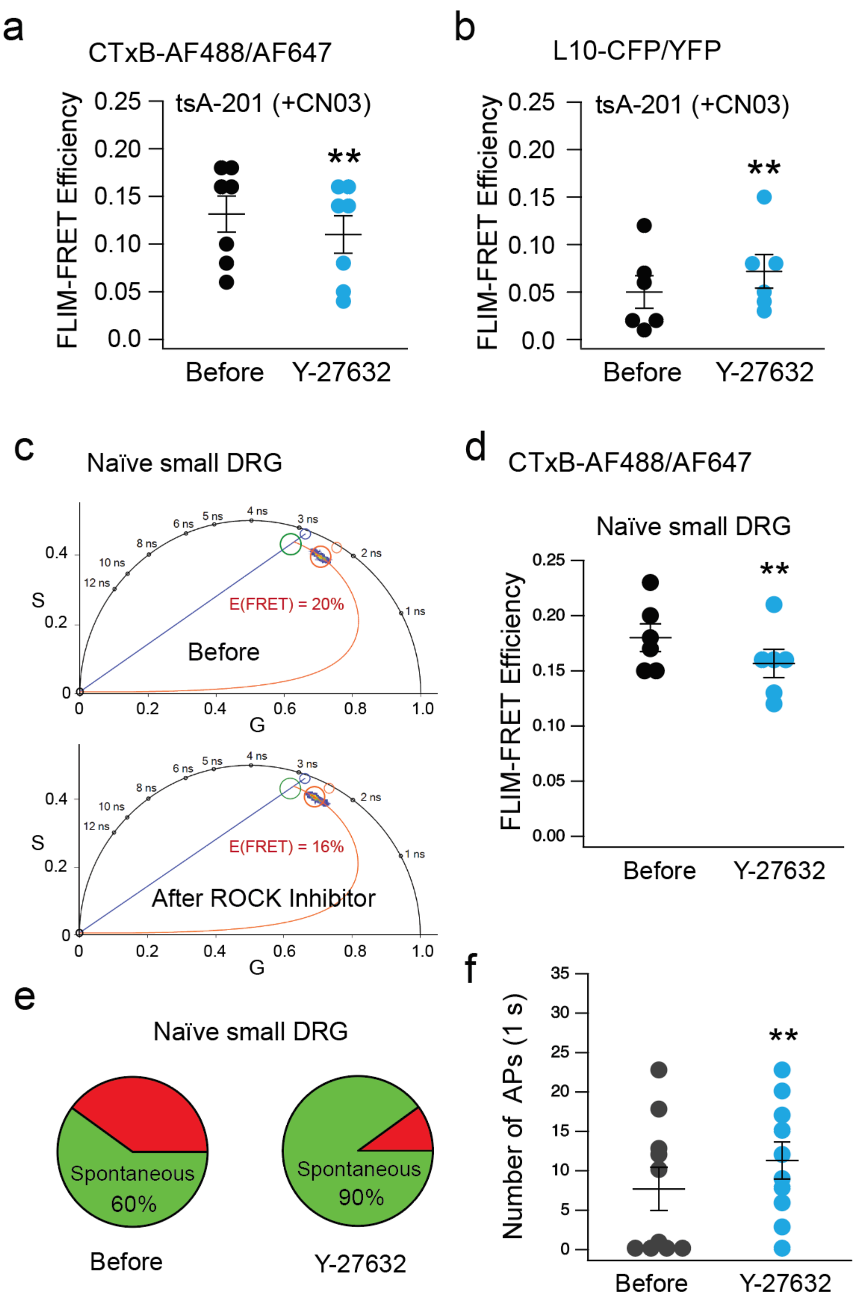


**Fig. S9. Effects of ROCK inhibition on membrane properties of naïve small DRG neurons. a-b.** Summary of the same-cell experiments showing the effects of ROCK inhibitor (20 μM Y-27632) on the L10-based (panel a, n = 6, p = 9e-4) and CTxB-based (panel b, n = 7, p = 6e-6) FRET measured from tsA cells following the overnight CN03 treatment, mean ± s.e.m., paired t-test. **c.** Representative phasor plots of small naïve DRG neurons labeled with CTxB-AF488 and CTxB-AF647 before and after ROCK inhibitor Y-27632 (20 μM). **d**. Summary of the FRET efficiency change for the CTxB-based FRET experiments as shown in panel c. Mean ± s.e.m., n = 6, p = 3e-3, paired t-test. **e**. Pie charts illustrating the fraction of small naïve DRG neurons that fired spontaneous APs under paired same-cell control conditions versus after the same acute ~ 5 min ROCK inhibitor treatment, n = 10 cells (independent dataset from those in Fig. 3c). **f**. Summary of spontaneous action potential firing in small naïve DRG neurons before and after acute Y-27632 treatment, n = 10, p = 6e-3, two-sided paired t-test.

**Supplementary Fig 10**

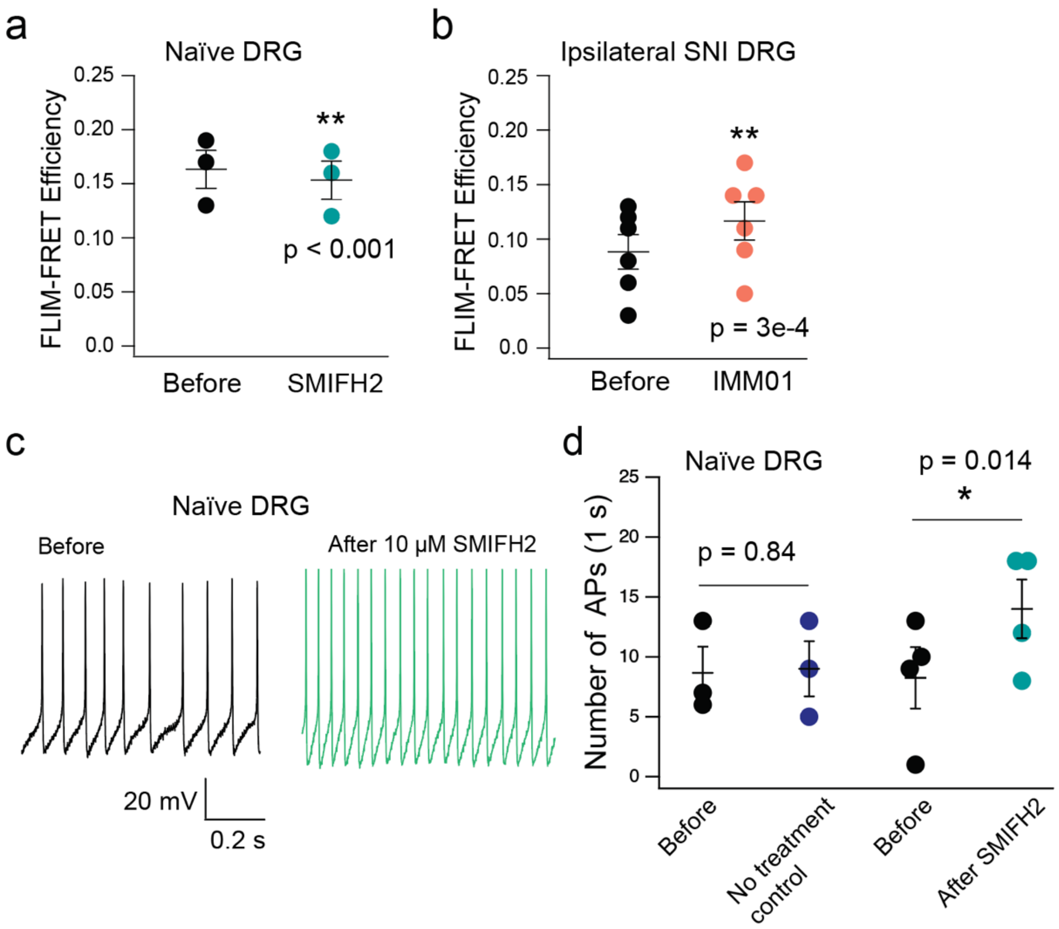


**Fig. S10. Effect of mDia activation and inhibition on small DRG neurons.**

**a-b**, Summary of the effects of acute SMIFH2 treatment on CTxB-based FRET in naïve small DRG neurons (a, n = 3, mean ± s.e.m.) and of IMM01 treatment on CTxB-based FRET in ipsilateral small DRG neurons following SNI (b, n = 6, mean ± s.e.m.), two-sided paired t-test. **c**, Representative recordings of spontaneous action potential firing from naïve small DRG neurons before and after acute SMIFH2 treatment. **d**, Summary of spontaneous action potential firing before and after acute SMIFH2 treatment in naïve DRG neurons, as well as the time control with no treatment. Data are mean ± s.e.m.; n = 4; paired two-tailed t-test, *p < 0.05, **p < 0.01.

**Supplementary Fig 11**

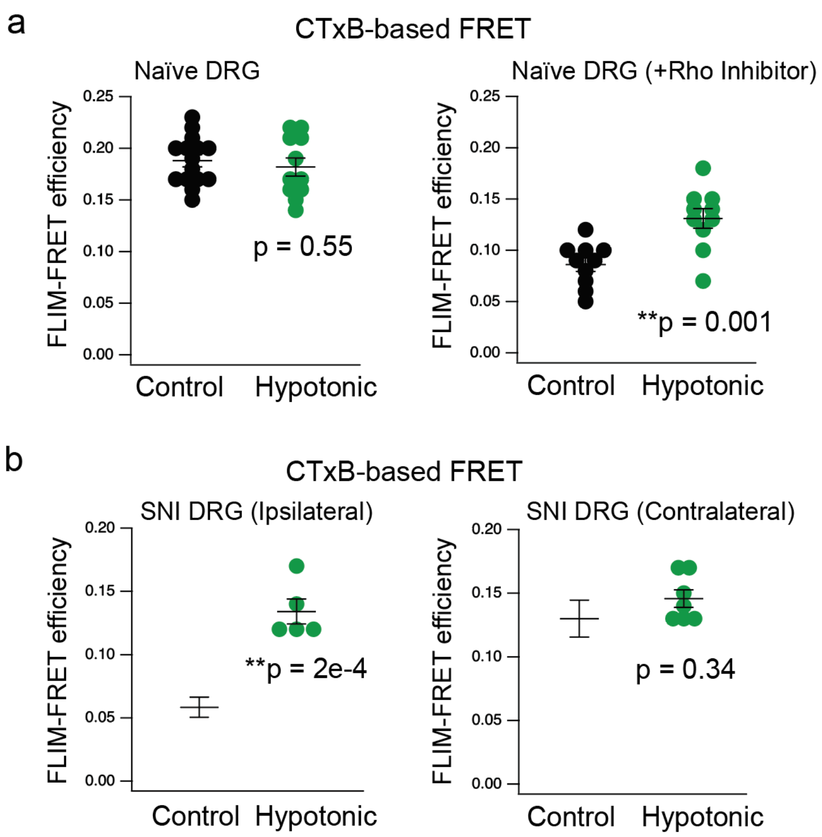


**Fig. S11. Hypotonic swelling increases CTxB-based FRET in small DRG neurons.**
**a**. Effects of acute (~3 min) hypotonic treatment at 154 mOsm on FLIM-FRET efficiency of naïve DRG neurons with or without overnight application of Rho inhibitor (exoenzyme C3 transferase).
**b**. Effects of acute hypotonic treatment on FLIM-FRET efficiency of ipsilateral and contralateral SNI pain neurons ; n = 5 cells for ipsilateral and n = 7 for contralateral SNI DRG neurons under the hypotonic condition. Data in control conditions are the same data as in Fig. 3d. Mean ± s.e.m., *p < 0.05, **p < 0.01, two-sided t-test.

**Supplementary Fig 12**

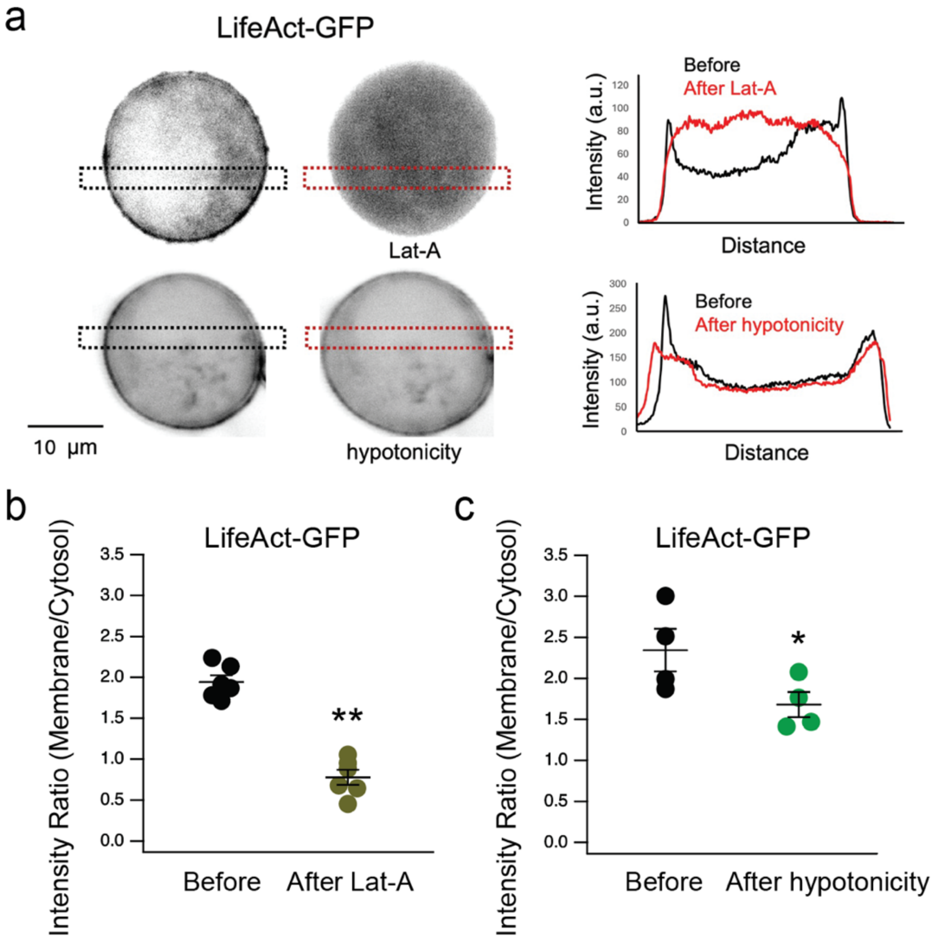


**Fig. S12. Effects of latrunculin A (Lat-A) and hypotonic treatments on cortical F-actin, visualized with LifeAct-GFP.**
**a**. Representative LifeAct-GFP fluorescence images during CN03 treatment and corresponding line-scan analyses across the dashed box, illustrating the reduction in membrane-associated F-actin signal. **b-c**. Summary of the membrane-to-cytosol fluorescence intensity ratio of LifeAct-GFP during acute treatment with Lat-A (n = 6, p = 5e-4) (b) or hypotonic solution (1:1 mix of the pre-made isotonic and hypotonic solutions) at 240 mOsm (n = 4, p = 0.043) (c), mean ± s.e.m., two-sided paired t-test.

**Table S1. Flipper-TR lifetimes* ($\tau_\phi$ and $\tau_m$) in small DRG neurons.**

| Cell type | Condition | $\tau_\phi$ (ns) | $\tau_m$ (ns) |
|---|---|---|---|
| Naïve Neuron | Untreated (n = 12) | 4.61 ± 0.04 | 4.72 ± 0.04 |
| | Rho Inhibitor (n = 11) | 4.45 ± 0.04 | 4.56 ± 0.02 |
| | Rho Activator (n = 12) | 4.66 ± 0.03 | 4.81 ± 0.04 |
| | BSA/2-BP (n = 11) | 4.32 ± 0.03 | 4.67 ± 0.03 |
| | BSA (n = 7) | 4.48 ± 0.04 | 4.74 ± 0.03 |
| | Lat-A (n = 9) | 4.57 ± 0.04 | 4.89 ± 0.05 |
| | BSA/Palmitate (n = 7) | 4.68 ± 0.02 | 4.78 ± 0.03 |
| Contralateral SNI Neuron | Untreated (n = 10) | 4.47 ± 0.03 | 4.87 ± 0.03 |
| | Rho Activator (n = 11) | 4.55 ± 0.03 | 4.90 ± 0.05 |
| | BSA/2-BP (n = 11) | 4.33 ± 0.03 | 4.74 ± 0.03 |
| | Lat-A (n = 10) | 4.72 ± 0.04 | 5.01 ± 0.08 |
| Ipsilateral SNI Neuron | Untreated (n = 13) | 4.36 ± 0.02 | 4.73 ± 0.03 |
| | Rho Activator (n = 12) | 4.58 ± 0.04 | 4.84 ± 0.04 |
| | BSA/2-BP (n = 10) | 4.40 ± 0.03 | 4.75 ± 0.05 |
| | Rho Activator + BSA/2-BP (n = 8) | 4.57 ± 0.02 | 4.84 ± 0.07 |
| | Lat-A (n = 11) | 4.66 ± 0.03 | 4.80 ± 0.05 |
| | Rho Activator + Lat-A (n = 14) | 4.62 ± 0.04 | 4.80 ± 0.03 |

*$\tau_\phi$ is preferred over $\tau_m$ for comparisons across experiments because it is less affected by background and noise, making it a more robust indicator of true fluorescence lifetime changes.
